# Hypomorphic mutation of the mouse Huntington’s disease gene orthologue

**DOI:** 10.1101/444059

**Authors:** Vidya Murthy, Toma Tebaldi, Toshimi Yoshida, Serkan Erdin, Teresa Calzonetti, Ravi Vijayvargia, Takshashila Tripathi, Emanuela Kerschbamer, Ihn Sik Seong, Alessandro Quattrone, Michael E. Talkowski, James F. Gusella, Katia Georgopoulos, Marcy E. MacDonald, Marta Biagioli

**Author notes:** These authors contributed equally to this work (Vidya Murthy, Toma Tebaldi). Corresponding author: Marta Biagioli, Ph.D., NeuroEpigenetics laboratory, Centre for Integrative Biology (CIBIO) University of Trento, Via Sommarive 938123 Povo (TN) - Italy.

## Abstract

Rare individuals with hypomorphic inactivating mutations in the Huntington’s Disease (HD) gene (*HTT*), identified by CAG repeat expansion in the eponymous neurodegenerative disorder, exhibit variable abnormalities that imply *HTT* essential roles during organ development. Here we report phenotypes produced when increasingly severe hypomorphic mutations in *Htt*, the murine *HTT* orthologue (in *Hdh*^neoQ20^, *Hdh*^neoQ50^, *Hdh*^neoQ111^ mice), were placed over a null allele (*Hdh*^ex4/5^). The most severe hypomorphic allele failed to rescue null lethality at gastrulation, while the intermediate alleles yielded perinatal lethality and a variety of fetal abnormalities affecting body size, skin, skeletal and ear formation, and transient defects in hematopoiesis. Comparative molecular analysis of wild-type and *Htt*-null retinoic acid-differentiated cells revealed gene network dysregulation associated with organ development and proposed polycomb repressive complexes and miRNAs as molecular mediators. Together these findings demonstrate that the HD gene acts both pre- and post-gastrulation and possibly suggest pleiotropic consequences of *HTT*-lowering therapeutic strategies.

**Author Summary:** The *HTT* gene product mutated in Huntington’s Disease (HD) has essential roles during normal organism development, however, still not fully predictable are the functional consequences of its partial inactivation. Our genetic study provides a comprehensive effects’ description of progressively stronger suppression of *Htt* gene, the murine *HTT* counterpart. The most severe *Htt* reduction leads to embryo lethality, while intermediate *Htt* dosages yield a variety of developmental abnormalities affecting body size, skin, skeletal and ear formation, and hematopoiesis. Complementing molecular analysis in differentiating cells depleted of a functional *Htt* gene further elucidates genes’ networks dysregulated during organ development and proposes chromatin regulators and short non-coding RNAs as key molecular mediators. Together these findings demonstrate that the HD gene acts both at early and later stages of development, thus possibly suggesting long-term consequences associated to the newest HD therapeutic strategies aimed at lowering the *HTT* gene product.

## Introduction

Huntington’s Disease (HD) is a dominantly inherited neurodegenerative disorder characterized by motor, cognitive and behavioral signs, generally of mid-life onset [1]. HD is caused by an unstable CAG trinucleotide repeat expansion in the 4p16.3 gene *HTT* (previously *HD*) [2]. The size of the expanded repeat is strongly correlated with age at onset but genetic variants at other loci, including DNA maintenance genes involved in somatic repeat instability, can modify the rate of pathogenesis [3].

Though knowledge of the pathogenic rate driver is emerging from HD genetic studies, it is not yet evident which aspect of the *HTT* expansion mutation is harmful to the cellular targets whose dysfunction or demise contributes to the disorder. The observation that CAG repeat expansions at different unrelated genes yield distinct neurological disorders argues that the harmful entity may be at the level of the *HTT*-encoded protein (huntingtin), through some opportunity afforded by the mutant protein’s normal function [4]. Consistent with this hypothesis, HD CAG expansion homozygotes exhibit onset similar to HD heterozygotes [5] and *HTT* inactivating mutations do not produce HD. Instead, rare humans with a single functional *HTT* copy [6], and mice with a single functional *HTT* orthologue (*Htt*, previously *Hdh*), whether a wild-type or a CAG repeat knock-in allele [7-9], are unremarkable. By contrast, complete inactivation of both alleles is early embryonic lethal [7,10,11].

The essential role played by *HTT* early in development is hypothesized, from genetic studies in the mouse, to intersect with chromatin regulation, influencing for example, the histone methyltransferase polycomb repressive complex 2 (PRC2) [12-14], although the lethality of null alleles has hampered studies later in development [7]. However, a recent report of a family segregating two rare, apparently incompletely inactivating *HTT* mutations [15,16], emphatically demonstrates that *HTT* is also essential for proper development of the brain and possibly other organs, as compound heterozygote children exhibit variable neurodevelopmental features, including motor disturbance, hypotonia, abnormal cranial circumference and small stature, as well as early death (OMIM: 617435) [16]. These observations are generally consistent with findings from genetic studies with members of a hypomorphic allelic series of flox-pGKneo-in CAG repeat knock-in mouse lines. *Hdh*^neoQ20^, *Hdh*^neoQ50^ and *Hdh*^neoQ111^ mice carry a mild, intermediate and severe hypomorphic neo-in allele, respectively [8,9,17]. Compound heterozygotes with combinations of these alleles, as well as homozygotes, exhibit a number of evident, variable abnormalities, involving brain development, movement deficits, decreased body size and reduced survival [9,17].

We have systematically evaluated each of the neo-in CAG repeat knock-in alleles for the ability to support proper development when placed over the same *Hdh*^ex4/5^ null allele in order to assess the extent to which dosage of the HD gene mouse orthologue, below a single functional copy, is essential for normal development and to uncover processes sensitive to gene dosage. Our findings, augmented by molecular analysis of *Hdh*^ex4/5^ embryonic stem cells (ESC) and retinoic acid differentiated cells, indicate that in addition to the brain, the proper development of several other organ systems requires huntingtin, as revealed by different developmental blocks that are by-passed at different dosages. Our data also highlight potentially responsible regulatory factors and networks that are sensitive to inactivating mutation of *Htt*.

## Results

### *Hdh*^neoQ20^, *Hdh*^neoQ50^, *Hdh*^neoQ111^ hypomorphic mutation and *Hdh*^ex4/5^ null rescue

*Hdh*^neoQ20^, *Hdh*^neoQ50^ and *Hdh*^neoQ111^ mice were bred to *Hdh*^ex4/5^ mice, with a targeted null allele [7], and their progeny were genotyped at weaning and at earlier stages, to assess the ability of each hypomorphic allele to rescue the early embryonic lethality at ~E7.5-8.0 imposed by complete gene inactivation. The results, summarized in Table 1, indicate the expected Mendelian ratio for progeny with wild-type alleles, at all ages, but revealed lethality for hypomorph mutation hemizygotes (with the *Hdh*^ex4/5^ null allele) beginning at progressively earlier stages with increased severity of inactivation. The *Hdh*^neoQ20^/*Hdh*^ex4/5^ diplotype was recovered at the expected Mendelian ratio at E14.5 and embryos appeared normal, while the *Hdh*^neoQ50/ex4/5^ was recovered at the expected Mendelian ratio at E10-12.5 but the embryos were smaller than wild-type. Finally, the *Hdh*^neoQ111^/*Hdh*^ex4/5^ diplotype was seen at E8.5, though all embryos had the abnormal sock-like appearance of *Hdh*^ex4/5^/*Hdh*^ex4/5^ null embryos just post-gastrulation, lacking head-folds (Figure 1A). *Hdh*^neoQ111^/*Hdh*^ex4/5^ embryos (non-resorbed) were not observed at later stages. *Hdh*^neoQ20^/*Hdh*^ex4/5^ and *Hdh*^neoQ50^/*Hdh*^ex4/5^ survivors were recovered at later stages but these died perinatally and at birth, respectively, displaying abnormalities that increased in severity with the gradient of inactivation *Hdh*^neoQ20^/*Hdh*^ex4/5^ < *Hdh*^neoQ50^/*Hdh*^ex4/5^: dome shaped cranium-to-overt exencephaly and mild-to-robustly decreased height and weight (Figure 1A-E).

**Table 1.** Summary of embryos recovered from matings of hypomorphic mutation mice with *Hdh*^ex4/5^/+ mice.

**A) *Hdh*<sup>neoQ20</sup>/+ X *Hdh*<sup>ex4/5</sup>/+ matings**
|  |  | WT |  | *Hdh*<sup>neoQ20</sup>/+ |  | *Hdh*<sup>ex4/5</sup>/+ |  | *Hdh*<sup>neoQ20</sup>/*Hdh*<sup>ex4/5</sup> |  |
| --- | --- | --- | --- | --- | --- | --- | --- | --- | --- |
| Age<br>(d.p.c) | Total<br>number<br>(Litters) | Normal<br>(%) | Abnormal<br>(%) | Normal<br>(%) | Abnormal<br>(%) | Normal<br>(%) | Abnormal<br>(%) | Normal<br>(%) | Abnormal<br>(%) |
| P0<br>Birth | 13<br>(1) | 5/5<br>(100) | 0/5<br>(0) | 3/3<br>(100) | 0/0<br>(0) | 4/4<br>(100) | 0/4<br>(0) | 0/1<br>(0) | 1/1<br>(100)* |
| E18.5 | 82<br>(6) | 17/17<br>(100) | 0/17<br>(0) | 23/23<br>(100) | 0/23<br>(0) | 20/20<br>(100) | 0/20<br>(0) | 0/22<br>(0) | 22/22<br>(100)** |
| E14.5 | 13<br>(1) | 5/5<br>(100) | 0/5<br>(0) | 2/2<br>(100) | 0/2<br>(0) | 3/3<br>(100) | 0/3<br>(0) | 3/3<br>(100) | 0/3<br>(0) |
| E11.5 | 22<br>(2) | 8/8<br>(100) | 0/8<br>(0) | 8/8<br>(100) | 0/8<br>(0) | 4/4<br>(100) | 0/4<br>(0) | 2/2<br>(100) | 0/2<br>(0) |

**\*1 small with mild dome head; \*\*17 small with mild dome head, 1 aborted, 4 resorbed** **B) *Hdh*<sup>neoQ50</sup>/+ X *Hdh*<sup>ex4/5</sup>/+ matings**
|  |  | WT |  | *Hdh*<sup>neoQ50</sup>/+ |  | *Hdh*<sup>ex4/5</sup>/+ |  | *Hdh*<sup>neoQ50</sup>/*Hdh*<sup>ex4/5</sup> |  |
| --- | --- | --- | --- | --- | --- | --- | --- | --- | --- |
| Age<br>(d.p.c) | Total<br>number<br>(Litters) | Normal<br>(%) | Abnormal<br>(%) | Normal<br>(%) | Abnormal<br>(%) | Normal<br>(%) | Abnormal<br>(%) | Normal<br>(%) | Abnormal<br>(%) |
| P0<br>Birth | 16<br>(2) | 3/3<br>(100) | 0/3<br>(0) | 10/10<br>(100) | 0/10<br>(0) | 3/3<br>(100) | 0/3<br>(0) | 0/0<br>(0) | 0/0<br>(0) |
| E18.5 | 267<br>(20) | 81/81<br>(100) | 0/81<br>(0) | 71/71<br>(100) | 0/71<br>(0) | 56/56<br>(100) | 0/56<br>(0) | 0/59<br>(0) | 59/59<br>(100)* |
| E14.5/<br>E15.5 | 32<br>(3) | 6/6<br>(100) | 0/6<br>(0) | 11/11<br>(100) | 0/11<br>(0) | 6/6<br>(100) | 0/6<br>(0) | 0/9<br>(0) | 9/9<br>(100)** |
| E12.5/<br>E13 | 25<br>(2) | 8/8<br>(100) | 0/8<br>(0) | 6/6<br>(100) | 0/6<br>(0) | 4/4<br>(100) | 0/4<br>(0) | 0/7<br>(0) | 7/7<br>(100)*** |
| E10.5/<br>E11 | 24<br>(2) | 6/6<br>(100) | 0/6<br>(0) | 7/7<br>(100) | 0/7<br>(0) | 5/5<br>(100) | 0/5<br>(0) | 0/7<br>(0) | 6/6<br>(100)*** |

|  |  | WT |  | *Hdh*<sup>neoQ111</sup>/+ |  | *Hdh*<sup>ex4/5</sup>/+ |  | *Hdh*<sup>neoQ111</sup>/*Hdh*<sup>ex4/5</sup> |  |
| --- | --- | --- | --- | --- | --- | --- | --- | --- | --- |
| Age<br>(d.p.c) | Total<br>number<br>(Litters) | Normal<br>(%) | Abnormal<br>(%) | Normal<br>(%) | Abnormal<br>(%) | Normal<br>(%) | Abnormal<br>(%) | Normal<br>(%) | Abnormal<br>(%) |
| E18.5 | 30<br>(3) | 8/8<br>(100) | 0/8<br>(0) | 9/9<br>(100) | 0/9<br>(0) | 13/13<br>(100) | 0/13<br>(0) | 0/0<br>(0) | 0/0<br>(0) |
| E14.5 | 13<br>(1) | 4/4<br>(100) | 0/4<br>(0) | 3/3<br>(100) | 0/3<br>(0) | 2/2<br>(100) | 0/2<br>(0) | 0/4<br>(0) | 4/4<br>(100)* |
| E10.5 | 10<br>(1) | 5/5<br>(100) | 0/5<br>(0) | 4/4<br>(100) | 0/4<br>(0) | 1/1<br>(100) | 0/1<br>(0) | 0/0<br>(0) | 0/0<br>(0) |
| E9 | 12<br>(1) | 4/4<br>(100) | 0/4<br>(0) | 1/1<br>(100) | 0/1<br>(0) | 4/4<br>(100) | 0/4<br>(0) | 0/3<br>(0) | 3/3<br>(100)** |
| E8.5 | 37<br>(3) | 12/12<br>(100) | 0/12<br>(0) | 7/7<br>(100) | 0/7<br>(0) | 5/5<br>(100) | 0/5<br>(0) | 0/13<br>(0) | 13/13<br>(100)** |
| E7 | 13<br>(1) | 7/7<br>(100) | 0/7<br>(0) | 3/3<br>(100) | 0/3<br>(0) | 2/2<br>(100) | 0/2<br>(0) | 1/1<br>(100) | 0/1<br>(0) |

\* resorbed; \*\*delayed sock-like embryonic portion

**Figure 1.**
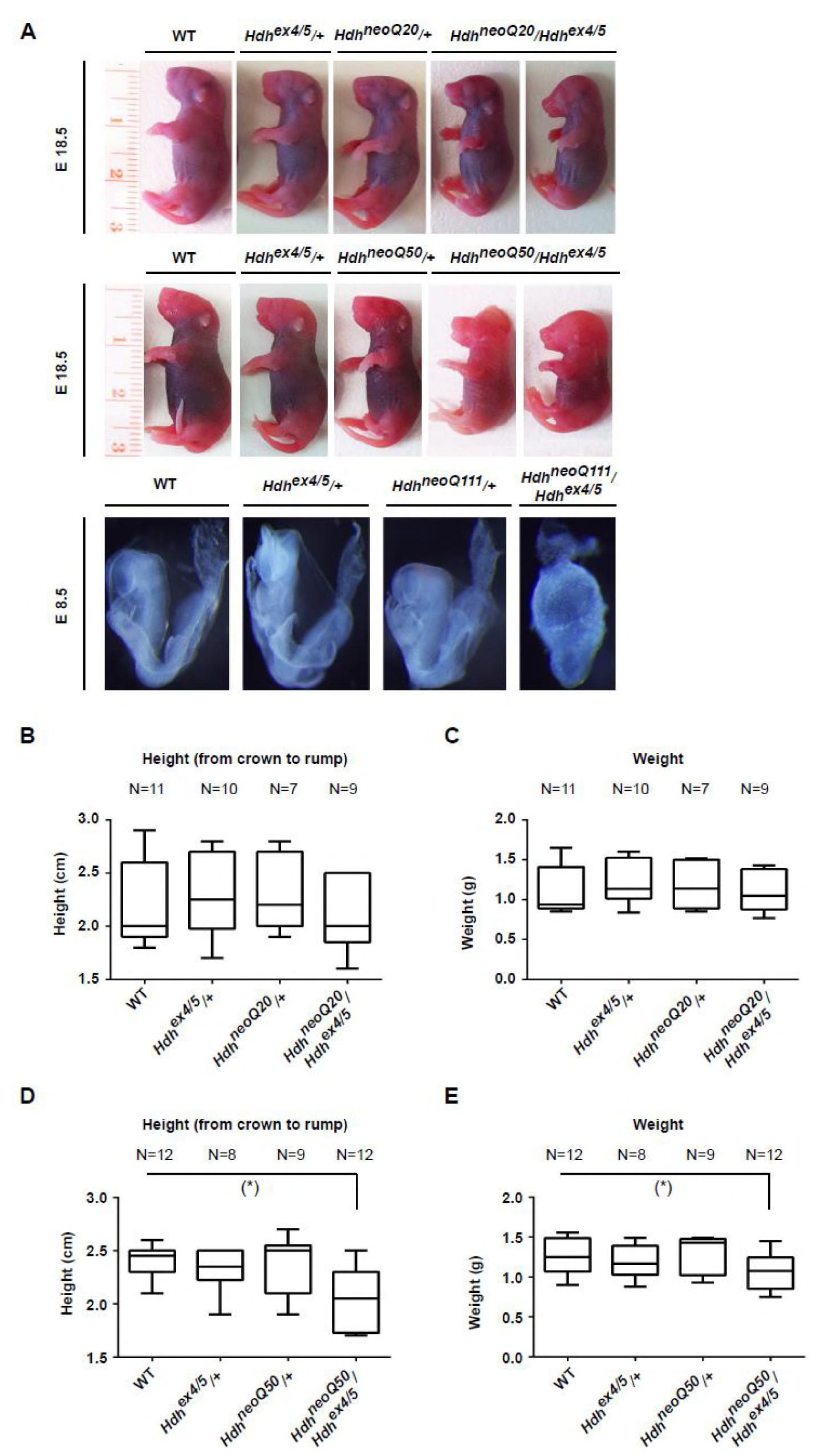
*Hdh*^neoQ20^/*Hdh*^ex4/5^, *Hdh*^neoQ50^/*Hdh*^ex4/5^ and *Hdh*^neoQ111^/*Hdh*^ex4/5^ hypomorphic embryos. **A)** Representative pictures of wild-type (WT), *Hdh*^ex4/5^/+, *Hdh*^neoQ20^/+, *Hdh*^neoQ50^/+, *Hdh*^neoQ111^/+, *Hdh*^neoQ20^/*Hdh*^ex4/5^, *Hdh*^neoQ50^/*Hdh*^ex4/5^ and *Hdh*^neoQ111^/*Hdh*^ex4/5^ hypomorphic embryos. The size and the shape of embryos with different genotypes is shown at E 18.5 for animals generated by crossing *Hdh*^neoQ20^/+ with *Hdh*^ex4/5^/+ mice and *Hdh*^neoQ50^/+ with *Hdh*^ex4/5^/+ mice. For embryos generated by crossing *Hdh*^neoQ111^/+ with *Hdh*^ex4/5^/+ mice, representative pictures of early stage animals were taken at E 8.5. See also Table 1 for full litters’ description and penetrance of the phenotypes. **B-C)** Bar graphs plot height (in cm) and weight (in grams - g) of different genotypic groups of animals generated by crossing *Hdh*^neoQ20^/+ with *Hdh*^ex4/5^/+ mice. The number (N) of animals for each group is indicated at the top of each bar. Error bars represent standard deviations from the mean. **D-E)** Bar graphs plot height (in cm) and weight (in grams - g) of different genotypic groups of animals generated by crossing *Hdh*^neoQ50^/+ with *Hdh*^ex4/5^/+ mice. The number (N) of animals for each group is indicated at the top of each bar. A significant reduction in size of *Hdh*^neoQ50^/*Hdh*^ex4/5^ hypomorphic embryos is noted (*). Error bars represent standard deviations from the mean.

### *Hdh*^neoQ20^/*Hdh*^ex4/5^ and *Hdh*^neoQ50^/*Hdh*^ex4/5^ have deficits in multiple organ systems

In previous studies of *Hdh*^ex4/5^/*Hdh*^ex4/5^ null embryos, embryoid bodies (EBs), as well as cultured ESC and neuronal cells, we demonstrated that a functional *Htt* allele is required for proper PRC2 activity, during genome-wide deposition of H3K27me3 epigenetic chromatin marks [12,13]. We therefore surveyed *Hdh*^neoQ20^/*Hdh*^ex4/5^ and *Hdh*^neoQ50^/*Hdh*^ex4/5^ embryos, mainly at E18.5, to evaluate a number of organ systems whose development is reported to require appropriate PRC2 activity and/or proper activity of its co-regulator polycomb repressive complex 1 (PRC1). Skeletal elements, ear development, skin barrier formation and fetal liver hematopoiesis were assessed.

### Skeleton

*Hdh*^neoQ20^/*Hdh*^ex4/5^ and *Hdh*^neoQ50^/*Hdh*^ex4/5^ embryos exhibited lumbar to sacral (L6 to the S1) vertebral transformation whose penetrance (20% and 30%, respectively) increased with the severity of inactivation. This abnormality was not observed in embryos with one wild-type allele (Figure 2A, Table 2). In addition, 50% of *Hdh*^neoQ20^/*Hdh*^ex4/5^ and 62.5% of *Hdh*^neoQ50^/*Hdh*^ex4/5^ embryos but none of the embryos with a wild-type allele displayed variable abnormalities of the sternum and xyphoid process (Figure 2B, Table 2), although the phenotypes were distinct. The *Hdh*^neoQ50^/*Hdh*^ex4/5^ xyphoid process was narrow, with reduced ossification, while the *Hdh*^neoQ20^/*Hdh*^ex4/5^ process was fenestrated. Furthermore, the tip of the *Hdh*^neoQ50^/*Hdh*^ex4/5^ sternum was bent inward, perhaps to accommodate a narrower thoracic cavity (Figure 2B, Table 2). The cervical vertebrae gap (C1 to C2 gap) was abnormally increased in 100% of *Hdh*^neoQ50^/*Hdh*^ex4/5^ embryos, perhaps secondary to the domed cranium and/or exencephaly (Figure 2C-D, Table 2). Variable hyoid bone and cartilage defects also increased with severity of the inactivating allele, such that 62% of *Hdh*^neoQ50^/*Hdh*^ex4/5^ and 10% of *Hdh*^neoQ20^/*Hdh*^ex4/5^ embryos had an abnormally short hyoid bone, greater horns and abnormal thyroid and cricoid cartilage morphology or spacing of the thyroid cartilage relative to the hyoid bone, respectively (Figure 2E, Table 2).

**Figure 2.**
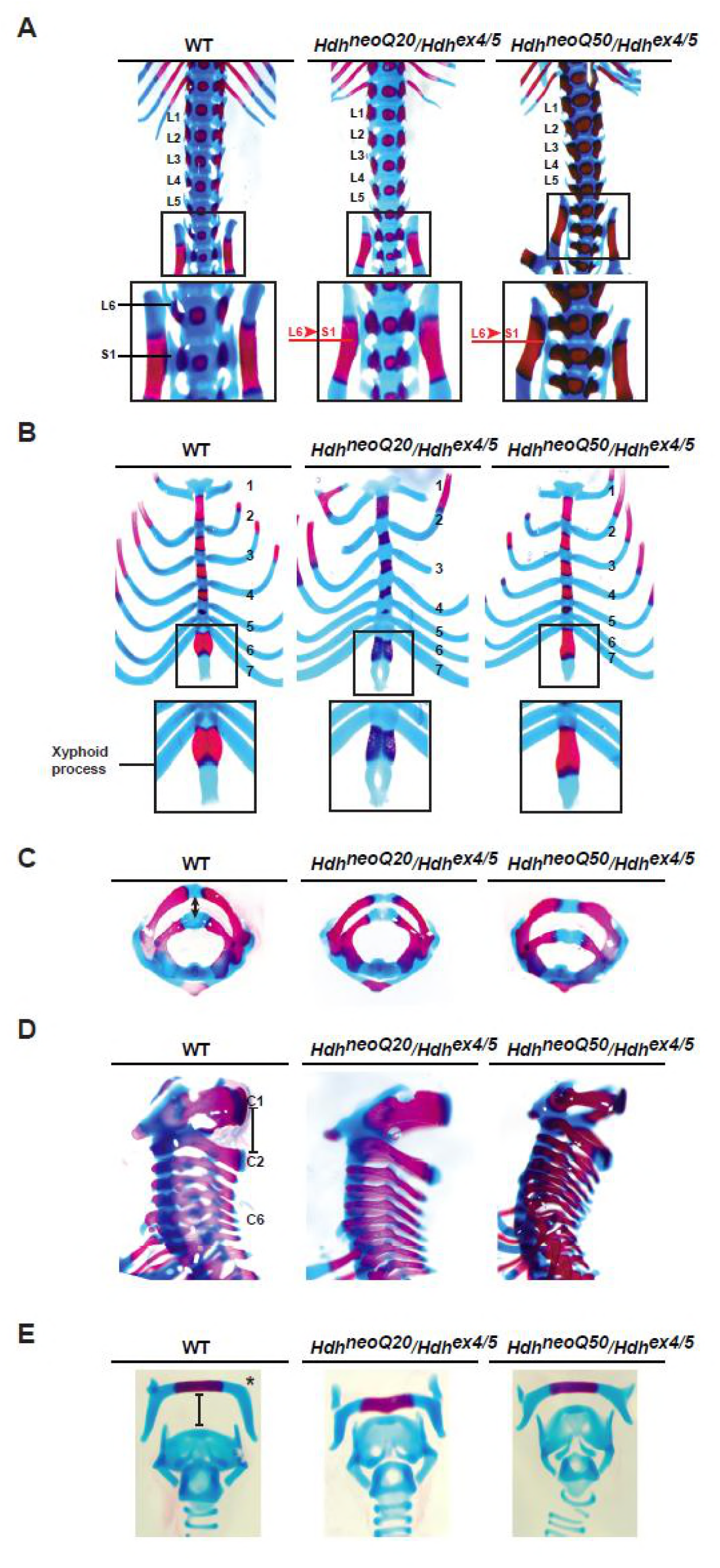
Skeletal defects in *Hdh*^neoQ20/*Hdh*ex4/5^ and *Hdh*^neoQ50/*Hdh*ex4/5^ hypomorphic embryos. Representative images of alcian blue and alizarin red-stained skeletal preparations from wild-type (WT), *Hdh*^neoQ20/*Hdh*ex4/5^, *Hdh*^neoQ50/*Hdh*ex4/5^, E 18.5 embryos showing (A) lumbar to sacral transformations (L6 to S1) in *Hdh*^neoQ20/*Hdh*ex4/5^ and *Hdh*^neoQ50/*Hdh*ex4/5^ embryos. The sacral area with the iliac bones is enlarged in the inset. L1 to L6 indicate lumbar vertebrae, while T13 refers to the 13th thoracic vertebra. Red typing depicts skeletal transformation; (B) Defects in the sternum and xyphoid process of *Hdh*^neoQ20/*Hdh*ex4/5^, *Hdh*^neoQ50/*Hdh*ex4/5^ mice, enlarged in the insets. The xyphoid process fenestration is clearly visible in *Hdh*^neoQ20/*Hdh*ex4/5^ embryos. Black numbers refer to thoracic ribs; (C-D) Cervical vertebrae defects in *Hdh*^neoQ20/*Hdh*ex4/5^ and *Hdh*^neoQ50/*Hdh*ex4/5^. Particularly, *Hdh*^neoQ50/*Hdh*ex4/5^ show an increase in the gap between C1 and C2 vertebrae, Horizontal views and vertical side views are shown in (C) and (D) respectively; E) Defects in the morphology [greater horns (*)] and spacing between hyoid bone and cartilage structures in the *Hdh*neoQ50/*Hdh*ex4/5 embryos. See also Table 2 for detailed description of the litters and quantitative analysis of the phenotypes.

**Table 2.** Summary of skeletal phenotypes in progeny of *Hdh*^neoQ20^/+ and *Hdh*^neoQ50^/+ X *Hdh*^ex4/5^/+ matings.

| Skeletal abnormalities | WT<br>Normal/<br>Abnormal<br>(%) | *Hdh*<sup>ex4/5/+</sup><br>Normal/<br>Abnormal<br>(%) | *Hdh*<sup>neoQ20/+</sup><br>Normal/<br>Abnormal<br>(%) | *Hdh*<sup>neoQ20/+</sup>/*Hdh*<sup>ex4/5</sup><br>Normal/<br>Abnormal<br>(%) | *Hdh*<sup>neoQ50/+</sup><br>Normal/<br>Abnormal<br>(%) | *Hdh*<sup>neoQ50/+</sup>/*Hdh*<sup>ex4/5</sup><br>Normal/<br>Abnormal<br>(%) |
| --- | --- | --- | --- | --- | --- | --- |
| **L6 to S1** | 0/8<br>(0) | 0/8<br>(0) | 0/6<br>(0) | 1/6*<br>(16.6) | 0/6<br>(0) | 2/8**<br>(25) |
| **Inverted**<br>**Sternum tip** | 0/8<br>(0) | 0/8<br>(0) | 0/6<br>(0) | 3/6<br>(50) | 0/6<br>(0) | 5/8<br>(62.5) |
| **Six**<br>**Vertebrosteral**<br>**ribs** | 0/8<br>(0) | 0/8<br>(0) | 0/6<br>(0) | 0/6<br>(0) | 0/6<br>(0) | 0/8<br>(0) |
| **Fenestrated**<br>**xiphoid process** | 0/8<br>(0) | 0/8<br>(0) | 0/6<br>(0) | 4/6<br>(66.6) | 0/6<br>(0) | 0/8<br>(0) |
| **T13 Aborted**<br>**ribs** | 0/8<br>(0) | 0/8<br>(0) | 0/6<br>(0) | 0/6<br>(0) | 0/6<br>(0) | 0/8<br>(0) |
| **C1 to C2 gap** | 0/8<br>(0) | 0/8<br>(0) | 0/6<br>(0) | 0/6<br>(0) | 0/6<br>(0) | 8/8<br>(100) |
| **C1 to C2** | 0/8<br>(0) | 0/8<br>(0) | 0/6<br>(0) | 0/6<br>(0) | 0/6<br>(0) | 0/8<br>(0) |
| **C7 to T1** | 0/8<br>(0) | 0/8<br>(0) | 0/6<br>(0) | 0/6<br>(0) | 0/6<br>(0) | 0/8<br>(0) |
| **Hyoid defect** | 0/8<br>(0) | 0/8<br>(0) | 0/6<br>(0) | 1/6<br>(16.6) | 0/6<br>(0) | 5/8<br>(62.5) |

\* Unilateral; \*\* Bilateral phenotypes.
\* Unilateral; \*\* Bilateral phenotypes.

### Ear

External ear and middle ear structures were also abnormal, increasing with the severity of the mutant allele: 100% of *Hdh*^neoQ50^/*Hdh*^ex4/5^ embryos had (bilateral) hypoplastic pinna and occasionally lack of the structure, while 33% of *Hdh*^neoQ20^/*Hdh*^ex4/5^ had (unilateral) hypoplastic pinna (Figure 3A-B, Table 3). Impact on middle ear structures further distinguished the inactivating alleles. Whereas all *Hdh*^neoQ20^/*Hdh*^ex4/5^ embryos had middle ear components, 75% of *Hdh*^neoQ50^/*Hdh*^ex4/5^ embryos lacked the tympanic ring, gonium, malleus, incus, and the third ossicle stapes, although Meckel’s cartilage was intact (Figure 3C, Table 3). The squamous bone was hypoplastic (Figure 3D, Table 3). For both inactivating mutations, inner ear structures were normal (data not shown). At E10.5, *Gsc* and *Hoxa2* expression in *Hdh*^neoQ50^/*Hdh*^ex4/5^ embryos was abnormal. *Gsc* mRNA, detected in the first and second branchial arches in wild-type embryos, was restricted to the first branchial arch. *Hoxa2* mRNA was reduced in intensity at the level of the branchial arches although intense staining was detected dorsally, suggesting a delayed or impaired migration of the neural-crest derived cells to colonize the arches, thereby possibly affecting the proper formation of ear structures (Figure 3E-F).

**Figure 3.**
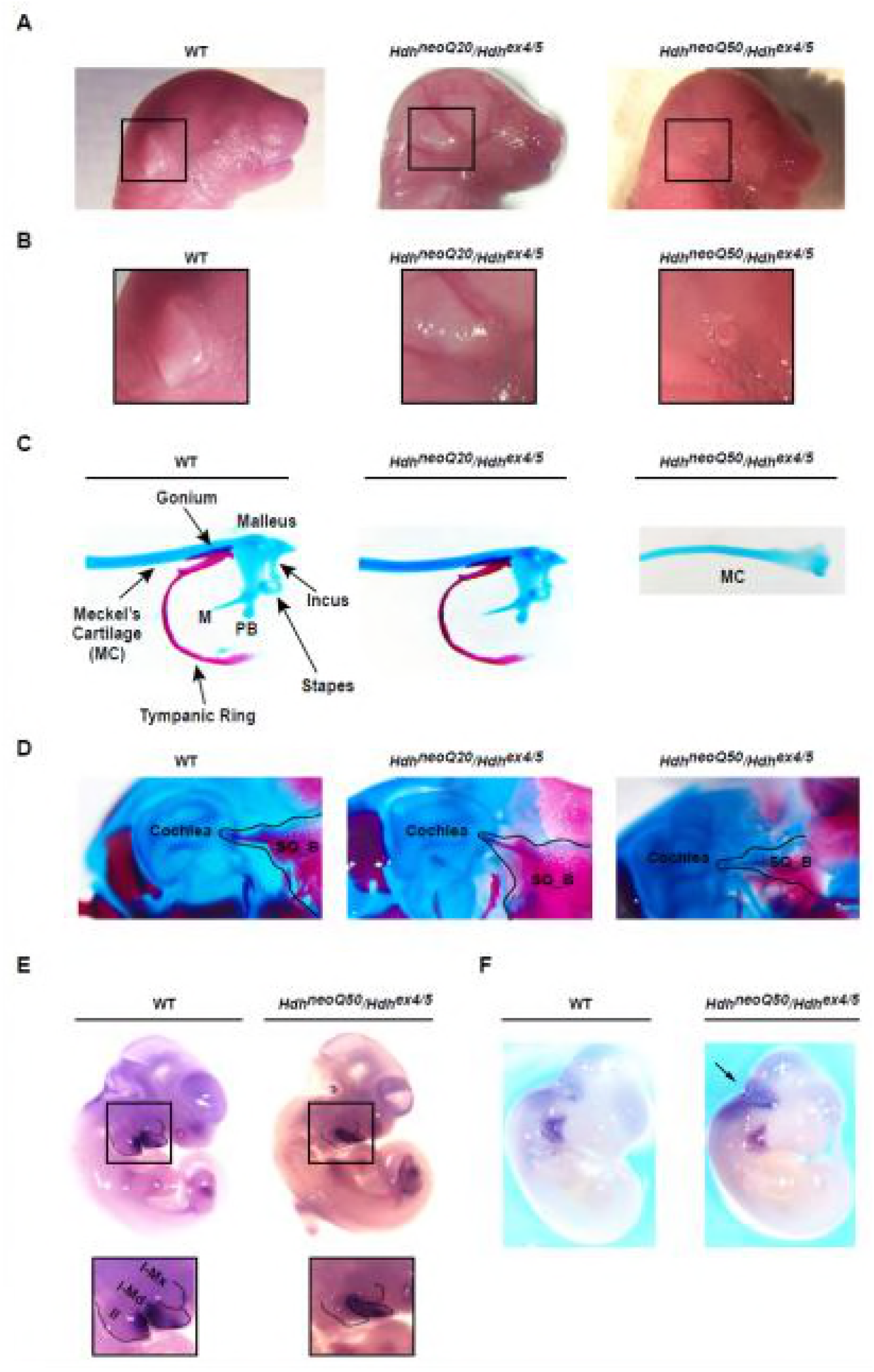
External and middle ear defects in *Hdh*^neoQ20^/*Hdh*^ex4/5^ and *Hdh*^neoQ50^/*Hdh*^ex4/5^ hypomorphic embryos. **A)** Representative images of wild-type (WT), *Hdh*^neoQ20^/*Hdh*^ex4/5^ and *Hdh*^neoQ50^/*Hdh*^ex4/5^ mice head describing altered morphology of external ear in *Hdh*^neoQ20^/*Hdh*^ex4/5^ and *Hdh*^neoQ50^/*Hdh*^ex4/5^ mice at E 18.5. **(B)** The region of the external ear is enlarged from (A) for each genotype, showing mis-shaped or lack of external ears for *Hdh*^neoQ20^/*Hdh*^ex4/5^ and *Hdh*^neoQ50^/*Hdh*^ex4/5^ hypomorphic embryos. **C)** Representative images of alcian blue and alizarin red-stained middle ear preparations from wild-type (WT), *Hdh*^neoQ20^/*Hdh*^ex4/5^, *Hdh*^neoQ50^/*Hdh*^ex4/5^E 18.5 embryos showing severely altered middle ossicles formation in *Hdh*^neoQ50^/*Hdh*^ex4/5^ hypomorfic embryos. Middle ear ossicles are named and indicated in the WT preparation. Abbreviations: M: manubrium, PB: processus brevis. In *Hdh*^neoQ50^/*Hdh*^ex4/5^ hypomorfic embryos all the middle ear ossicles were absent, only the Meckel’s Cartilage (MC) remained unaltered. **D)** Representative images of alcian blue and alizarin red-stained middle ear preparations from wild-type (WT), *Hdh*^neoQ20^/*Hdh*^ex4/5^, *Hdh*^neoQ50^/*Hdh*^ex4/5^E 18.5 embryos showing hypoplastic squamous bone (SQ_B) in *Hdh*^neoQ20^/*Hdh*^ex4/5^ and *Hdh*^neoQ50^/*Hdh*^ex4/5^ hypomorfic embryos. Black line delineates the squamosal bone. **E)** Whole mount *in situ* hybridization representative images showing altered *Gsc* gene localization in *Hdh*^neoQ50^/*Hdh*^ex4/5^ hypomorfic embryos at E11.5. The inset shows an enlarged view of the pharyngeal arches with *Gsc* transcript staining localized in the second and third arch in WT, but only localized in the first arch in *Hdh*^neoQ50^/*Hdh*^ex4/5^ embryos. Abbreviations: I-md, mandibular process of pharyngeal arch 1; I-mx, maxillary process of pharyngeal arch 1; II: pharyngeal arch 2. Dashed lines delineated different pharyngeal arches districts. **F)** Whole mount *in situ* hybridization representative images showing altered *Hoxa2* gene localization in *Hdh*^neoQ50^/*Hdh*^ex4/5^ hypomorphic embryos at E11.5. The arrows indicated ectopic expression of *Hoxa2* transcript in the dorsal area of *Hdh*^neoQ50^/*Hdh*^ex4/5^ embryos.

**Table 3.**
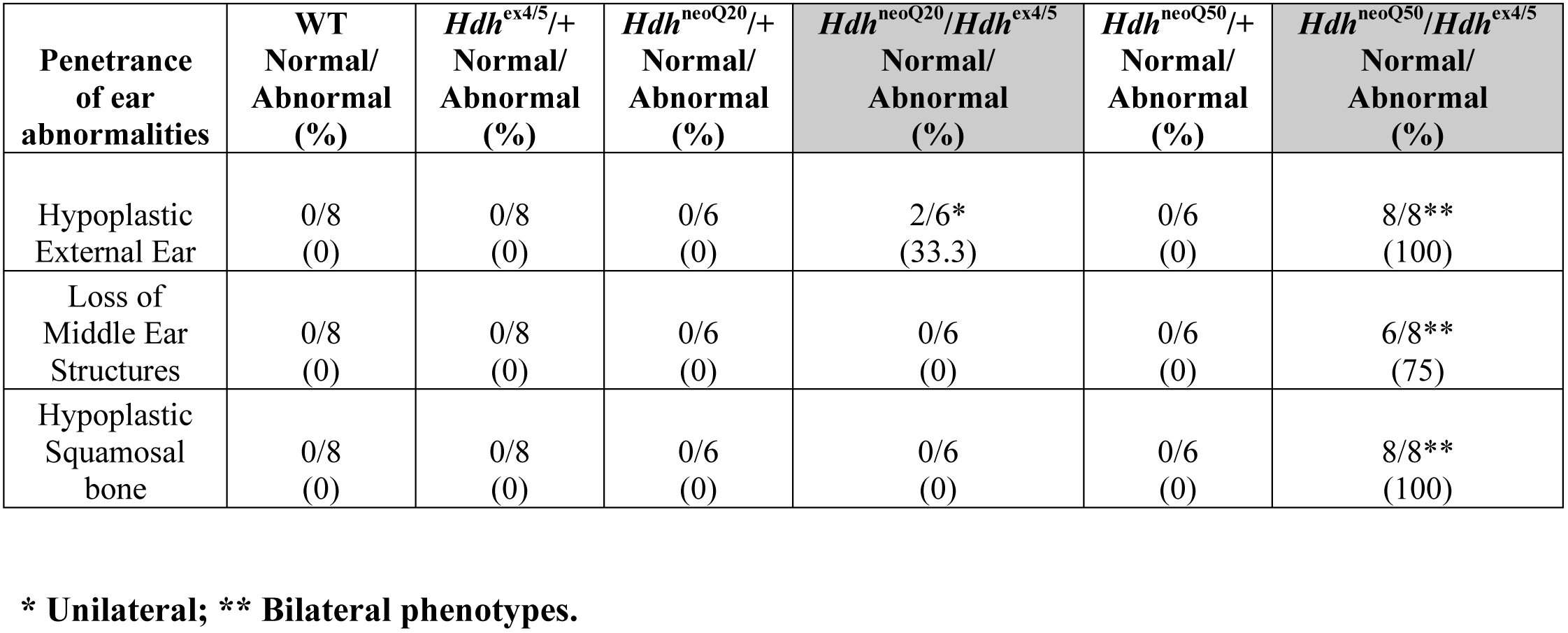
Summary of ear phenotypes in progeny of *Hdh*^neoQ20^/+ and *Hdh*^neoQ50^/+ X *Hdh*^ex4/5^/+ matings.

### Skin barrier

E18.5 *Hdh*^neoQ20^/*Hdh*^ex4/5^ embryos, like embryos with a wild-type allele, excluded toluidine blue dye, demonstrating proper formation of the cornified layer that serves as the skin barrier, but 100% of the *Hdh*^neoQ50^/*Hdh*^ex4/5^ hypomorphs excluded dye from the dorsal, but not the ventral epidermis (Figure 4A). The four stratified layers of the epidermis (basal, spinous, granular, cornified) were present (data not shown). However, immunoblot analysis indicated decreased levels of mature filaggrin (27 kDa, epidermal granular layer marker [18]), profilaggrin processing intermediates (50-70 kDa), and of loricrin (epidermal cornified layer marker) (Figure 4B). *Hdh*^neoQ50^/*Hdh*^ex4/5^ ventral epidermis also exhibited decreased PCNA staining (S-phase cell cycle marker) and increased TdT-mediated dUTP nick-end labeling (TUNEL)-positive apoptotic cells, compared to wild-type (Figure 4C-D), consistent with failed differentiation.

**Figure 4.**
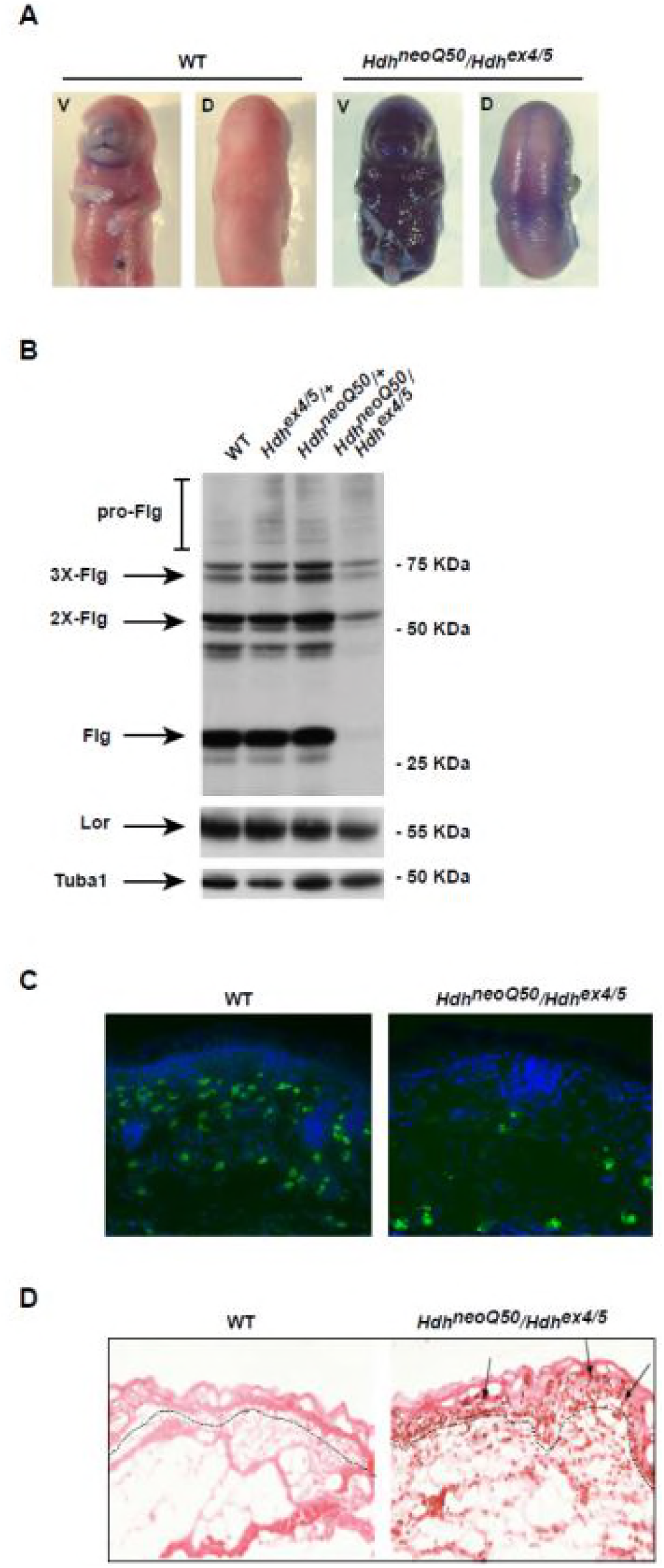
Skin defects in *Hdh*^neoQ20^/*Hdh*^ex4/5^ and *Hdh*^neoQ50^/*Hdh*^ex4/5^ hypomorphic embryos. Representative pictures of *Hdh*^neoQ50^/*Hdh*^ex4/5^ hypomorphic **(A)** E 18.5 mice following toluidine-blue dye exclusion assays. A functional epidermal barrier is shown in wild-type mice, while *Hdh*^neoQ50^/*Hdh*^ex4/5^ hypomorphs exhibited a compromised epidermal barrier on the ventral surface. **B)** Immunoblot analyses of wild-type (WT), *Hdh*^ex4/5^/+, *Hdh*^neoQ50^/+, *Hdh*^neoQ50^/*Hdh*^ex4/5^ embryos’ skin extracts. Pro-filaggrin (pro-Flg - 50, 70 KDa and higher molecular weight) and processed filaggrin (Flg-27 KDa) bands were detected in all samples, but they appered greatly reduced in *Hdh*^neoQ50^/*Hdh*^ex4/5^ embryos’ skin extracts. The bands of Loricrin (Lor) and alpha-tubulin (Tuba1) were detected unchanged in all samples. **C)** Representative images of cryostat-sectioned skin tissue, with DAPI stained nuclei to show reduced staining of the PCNA proliferation marker in *Hdh*^neoQ50^/*Hdh*^ex4/5^ hypomorphic embryos. Green, PCNA staining; Blue, DAPI-stained nuclei. Dashed lines depict the cornified basal epidermal layer. All images were taken using a fluorescent 20X objective. **D)** Representative pictures of cryostat-sectioned skin tissue stained with TUNEL assay to quantify apoptotic cell death (arrows) in *Hdh*^neoQ50^/*Hdh*^ex4/5^ hypomorphic embryos. All images were taken using a phase-contrast 20X objective.

### Hematopoiesis

At E14.5, *Hdh*^neoQ20^/*Hdh*^ex4/5^ embryos were unremarkable but *Hdh*^neoQ50^/*Hdh*^ex4/5^ embryos had small, pale livers, contrasting with abnormal vasculature and extensive blood inclusions in the head (brain and ventricles) reported previously [9] (Figure 5A, Suppl. Fig. 1). Flow cytometry (FACS) of fetal liver cells showed decreased absolute cell number and lack of erythropoiesis (CD71^hi^Ter119^neg-lo^ proerythroblasts to CD71^hi^Ter119^+^ basophilic erythroblasts to CD71^+^Ter119^+^ polychromatic erythroblasts to CD71^−^Ter119^+^ reticulocytes) in *Hdh*^neoQ50^/*Hdh*^ex4/5^ embryos (Figure 5B, C), concomitant with increased numbers of myeloid (Mac-1^+^Gr-1^−^) and B cell progenitors (B220^+^CD19^−^) (Figure 5B, C). Furthermore, the numbers of erythroid (CFU-E) and megakaryocyte (CFU-Mk) colony forming progenitors were decreased, along with a decrease in the total number of colony forming cells (Figure 5D), implying impaired expansion of HSCs/progenitor cells and defective erythropoiesis. The impairment was transient, as E18.5 *Hdh*^neoQ50^/*Hdh*^ex4/5^ livers did not show these hematopoietic deficits (Figure 5E-I). Analysis of thymus at this age also did not reveal deficits in the development of CD4^+^CD8^+^ double positive T cells suggesting normal T-lymphopoiesis, despite an apparently decreased number of thymic precursors (Figure 5E-I).

**Figure 5.**
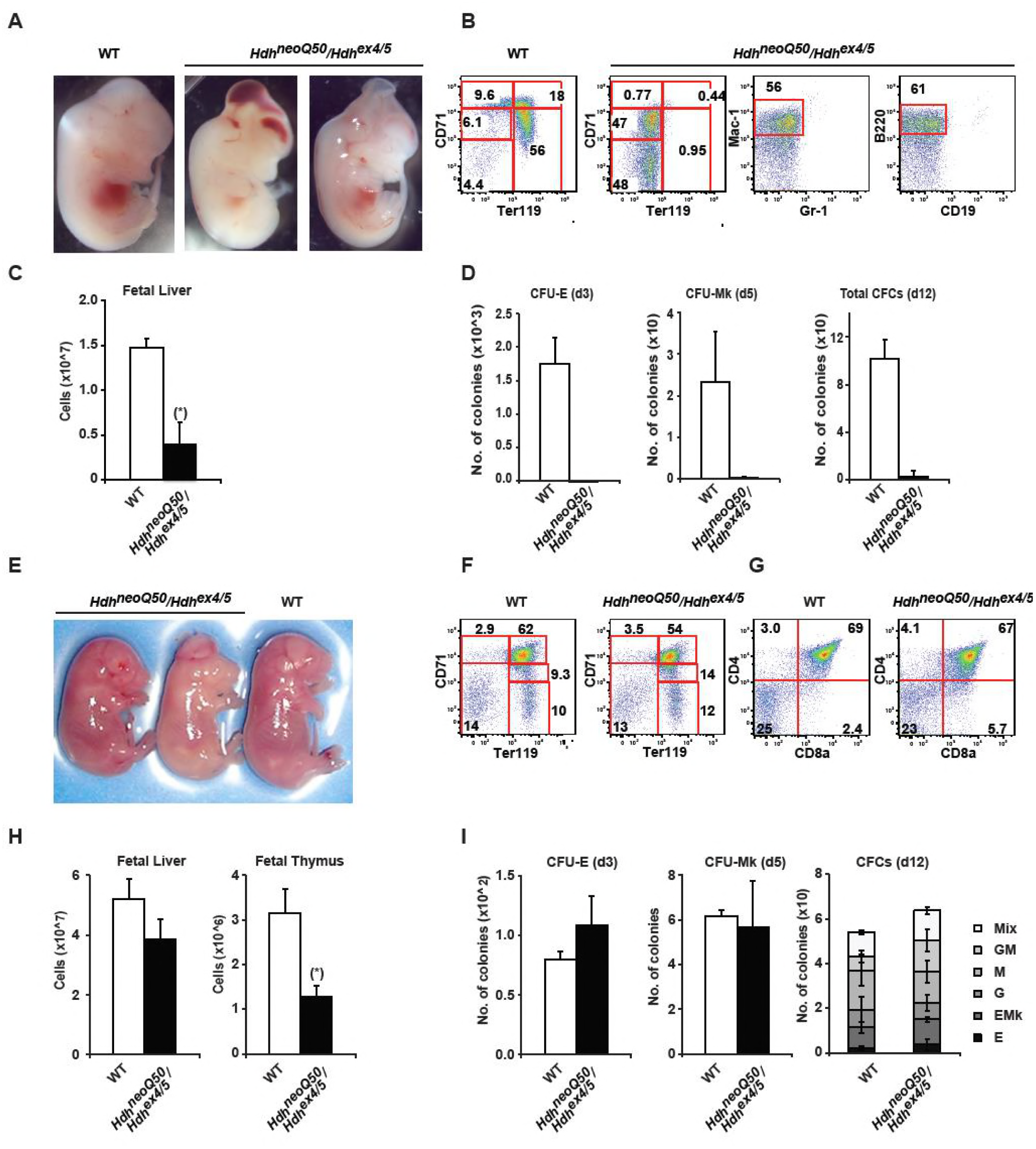
Hematopoietic defects in *Hdh*^neoQ50^/*Hdh*^ex4/5^ hypomorphic embryos. **(A, E)** Representative pictures of the appearance of fetal liver in wild-type (WT) and *Hdh*^neoQ50^/*Hdh*^ex4/5^ embryos displaying dome shaped cranium and exencephaly at E 14.5 and E 18.5. **(B, F, G**) Representative FACS profiles of fetal liver hematopoiesis in wild-type (WT) and *Hdh*^neoQ50^/*Hdh*^ex4/5^ embryos at E 14.5 and 18.5 respectively. Erythroblasts were gated into fractions defined by CD71 and Ter119 expression, displaying transient anemia at E 14.5, rescued by E 18.5. Myeloid cells and B cell progenitors were identified as Mac-1^+^Gr-1^−^ and B220^+^CD19^−^, respectively at E 14.5 Fetal liver T cells were stained with CD4 and CD8a cell surface markers. **(C, H)** Bar graph reporting the absolute cell numbers of whole fetal liver at E 14.5 and E 18.5 for wild-type (WT) and *Hdh*^neoQ50^/*Hdh*^ex4/5^ embryos, displaying transient defects in hematopoiesis at E 14.5. WT embryos N=5, *Hdh*^neoQ50^/*Hdh*^ex4/5^ embryos N= 3 at E 14. 5. WT embryos N=3, *Hdh*^neoQ50^/*Hdh*^ex4/5^ embryos N= 3 at E 18. 5. Error bars represent standard deviations from the mean. (*) indicates significant *P* value, *P*<0.01. **(D, I)** Bar graph reporting the results of colony assays on wild-type (WT) and *Hdh*^neoQ50^/*Hdh*^ex4/5^ fetal liver indicating a dramatic depletion of erythroid (CFU-E), Megakaryocytes (CFU-Mk) as well as the total colony forming cells in E 14.5 *Hdh*^neoQ50^/*Hdh*^ex4/5^ fetal liver. By E 18.5 the phenotype was completely rescued and no difference in erythroid (CFU-E and BFU-E (E)), megakaryocyte (CFU-Mk) myeloid (G, M, GM), erythroid-megakaryocyte (E/Mk) and multi-lineage mixed (Mix) colony forming cells was observed. Error bars represent standard deviations from the mean. d3, day3; d5, day5, d12, day12.

### *Hdh*^ex4/5/ex4/5^ developmental response to retinoic acid-induced differentiation

In comparisons of wild-type and *Hdh*^ex4/5/ex4/5^ null embryonic stem cells (ESC), at baseline and after Embryoid Bodies (EB) retinoic acid (RA)-induced differentiation, we have found previously that complete *Htt* inactivation impairs PRC2 activity [12,13] and alters genome-wide deposition of epigenetic chromatin marks [12,13]. Since RA-differentiation of ESC provides a culture system with which to study not only neurogenesis but organogenesis [19-21] more broadly, we utilized this paradigm to identify transcriptional regulators responsive to *Htt* dosage that may play a role in the incomplete hypomorph rescue of the *Hdh*^ex4/5/ex4/5^ null allele. Specifically, we performed differential gene expression analyses of genome-wide RNA and miRNA next generation sequencing data (Suppl. Table 1) generated from wild-type parental ESC, *Hdh*^ex4/5/ex4/5^ ESC and from wild-type and *Hdh*^ex4/5/ex4/5^ EB differentiated cells, treated with RA. The overall gene expression changes, upon RA-induced differentiation, were quantitatively and qualitatively similar for wild-type and *Hdh*^ex4/5/ex4/5^ cells: 79% of the several hundred changed RA-responsive miRNAs were shared (Suppl. Fig. 2, Suppl. File S1) and 86% of the more than 9,000 mRNA gene expression changes were shared (Suppl. Fig 2, Suppl. File S1). Functional annotation enrichment analysis (clusterProfiler) [22] highlighted genes involved in organogenesis in a large differentiation up-regulated cluster (Cluster 1, Suppl. Fig. 2E), associated with terms such as Skeleton, Ossification, Ear, Skin and Neuron development (Suppl. Fig. 2, Suppl. File S2), as well as genes involved in subcellular processes in a smaller down-regulated class (Cluster 2) (Suppl. Fig. 2E), with enriched terms such as DNA metabolic process, Ribosome biogenesis, Cell Cycle (Suppl. Fig. 2, Suppl. File S2). We then determined the impact of *Htt* nullizigosity on development, assessing the initial pluripotent stage (wild-type versus *Hdh*^ex4/5/ex4/5^ ESC) as well as the differentiated stage (wild-type versus *Hdh*^ex4/5/ex4/5^ RA-induced) (Suppl. Fig.3, Suppl. File S1). The results disclosed a modest number of miRNAs (ESC: 149 up: 95 down; RA: 70 up; 86 down) and mRNAs (ESC: 699 up: 461 down; RA 1314 up; 1784 down), whose expression was sensitive to *Htt* inactivation. These were largely specific to a given developmental stage (Suppl. Fig.3, Suppl. File S1), forming four non-overlapping gene sets (ESC-up, ESC-down, RA-up and RA-down) (Suppl. Fig. 3, Suppl. File S2). Gene Ontology and pathway enrichment analysis revealed highly significant enrichment of the *Htt*-inactivation sensitive RA-down gene set in developmental pathways related to organ system deficits observed in the hypomorphic mice: ‘skeletal system development’, ‘generation of neurons’, ‘blood vessel morphogenesis’, ‘blood vessel morphogenesis’, “skin development” and “ear development” (Suppl. Fig. 3).

In addition, we examined the interplay between RA-differentiation and *Htt*-inactivation, generating four gene expression sets: up_up, up_down, down_up and down_down, depending upon whether expression of a particular gene or miRNA was significantly up- or down-regulated by RA-differentiation and further was up- or down-regulated by *Hdh*^ex4/5^/*Hdh*^ex4/5^ null mutation, respectively (Fig. 6A-B, Suppl. File S1). Interestingly, miRNAs were evenly distributed across these classes (Suppl. File S1), while the mRNA expression changes were largely in the up_down class (Figure 6C). With these RA-differentiation-*Htt*-null gene sets we performed pathways analyses to assess membership in: 1) available Gene Ontology biological processes, 2) manually curated lists of genes relevant to organ development found to be abnormal in hypomorphic mice (Materials and Methods, Suppl. Table 2 and Suppl. File S1), and, 3) lists of genes expressed in specific brain cell types (210 neuron genes, 144 astrocyte genes, 61 oligodendrocyte genes [23], specifically to assess the hypothesis that *Htt*-null mutation alters the neuron-glia developmental switch [24]. The results of Gene Ontology analysis showed significant enrichment, mostly for ‘up_down but also the ‘up_up gene sets, highly associated with the terms ‘Organ Morphogenesis’, ‘Nervous System Development’, ‘Skeletal System Morphogenesis’, ‘Blood Vessel Morphogenesis’, ‘Ear Development’ and “Skin Development” (Fig. 6D, Suppl. File S2). Analysis with our manually-curated lists of genes involved in body weight (bodyweight), skin differentiation (skin), skeleton development (skeleton), middle-ear development (middle-ear) and hematopoiesis revealed significant enrichment scores for all of these hypomorph phenotype-related pathways, with the majority of the genes belonging to the ‘up-down’ class being up-regulated by RA-differentiation and down-regulated by *Htt*-null mutation (Fig. 7A-C). Furthermore, this class also contained most of the differentially regulated astrocyte and oligodendrocyte specific genes, whereas most of the neuron-specific genes were in the up-up class, up-regulated by RA differentiation and further up-regulated by *Htt*-null mutation (Fig. Suppl. 4, Suppl. File S1). This differential impact on genes expressed in neurons and in the major macroglial cell types strongly implied an effect on neurogenesis that, by analogy, implied effects of *Htt* loss of function mutation on progenitor stem cells.

**Figure 6.**
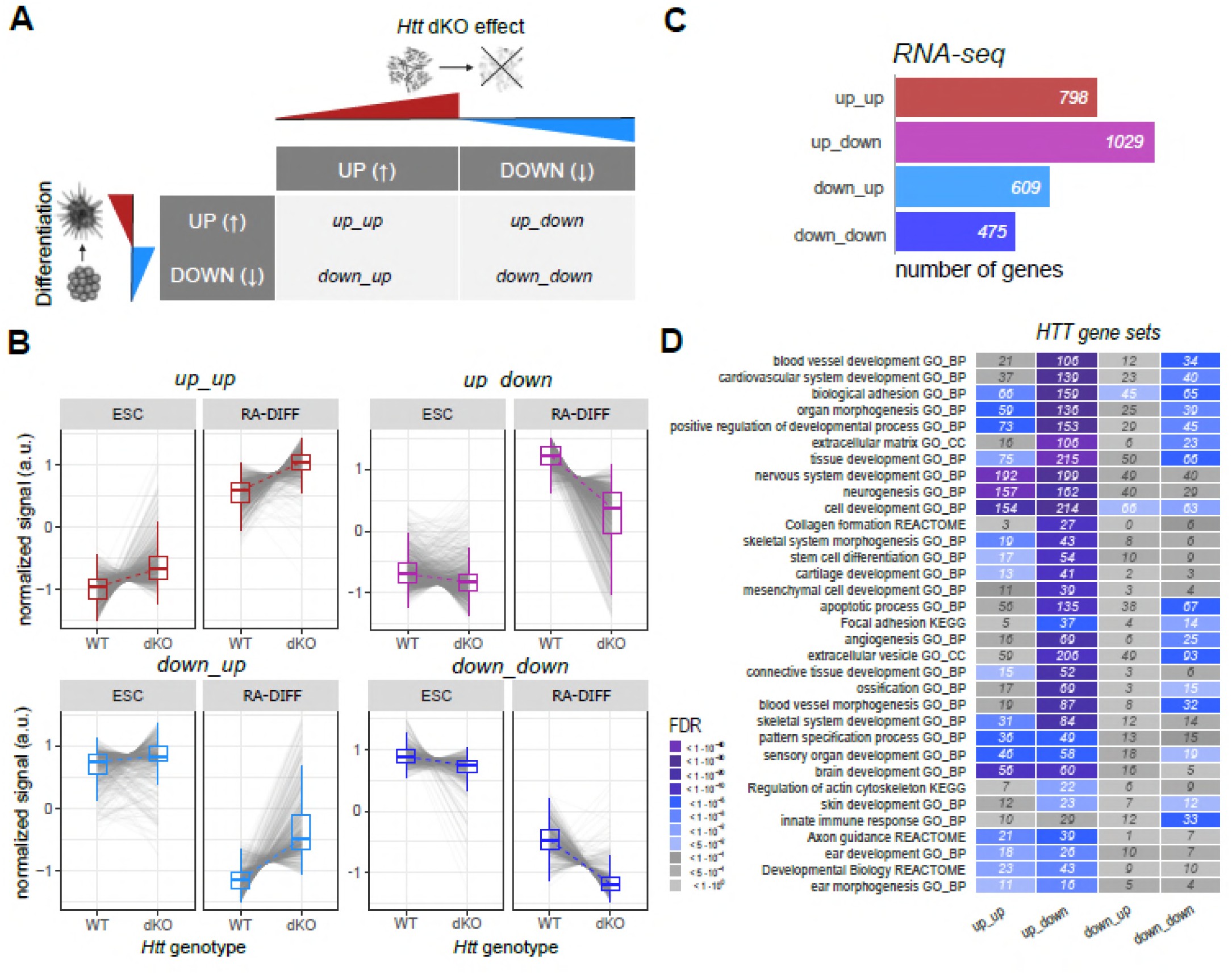
Classes of genes implicated in organ development whose expression is altered by *Htt*^ex4/5^ null. A) Schematic describing the criteria applied to define the 4 classes of genes sensitive to both *Hdh*^ex4/5^ null mutation and RA-induced differentiation. Up_up: genes up-regulated during differentiation and up-regulated by the *Htt*-null mutation; up_down: genes up-regulated during differentiation but down-regulated by *Htt*-null mutation; down_down: genes down-regulated during differentiation and further down-regulated by *Htt*-null mutation; down_up: genes down-regulated during differentiation, but upregulated in the context of *Htt*-null mutation. B) Expression trajectory plots describing variations in transcriptional levels across the two *Htt* genotypes (WT and dKO) and two developmental stages (ESC and RA-DIFF) of the genes belonging to the 4 categories described in A). C) Enrichment analysis of genes belonging to the 4 categories (Up_enhanced; up_counteracted; down_enhanced; down_counteracted, as described in A), highlighting the most enriched GO-terms [BP, Biological Process; CC, Cellular Component; MF; Molecular Function; KEGGs pathways and Reactome pathways]. Significance of enrichment is based on FDR values [color code as in the legend].

**Figure 7.**
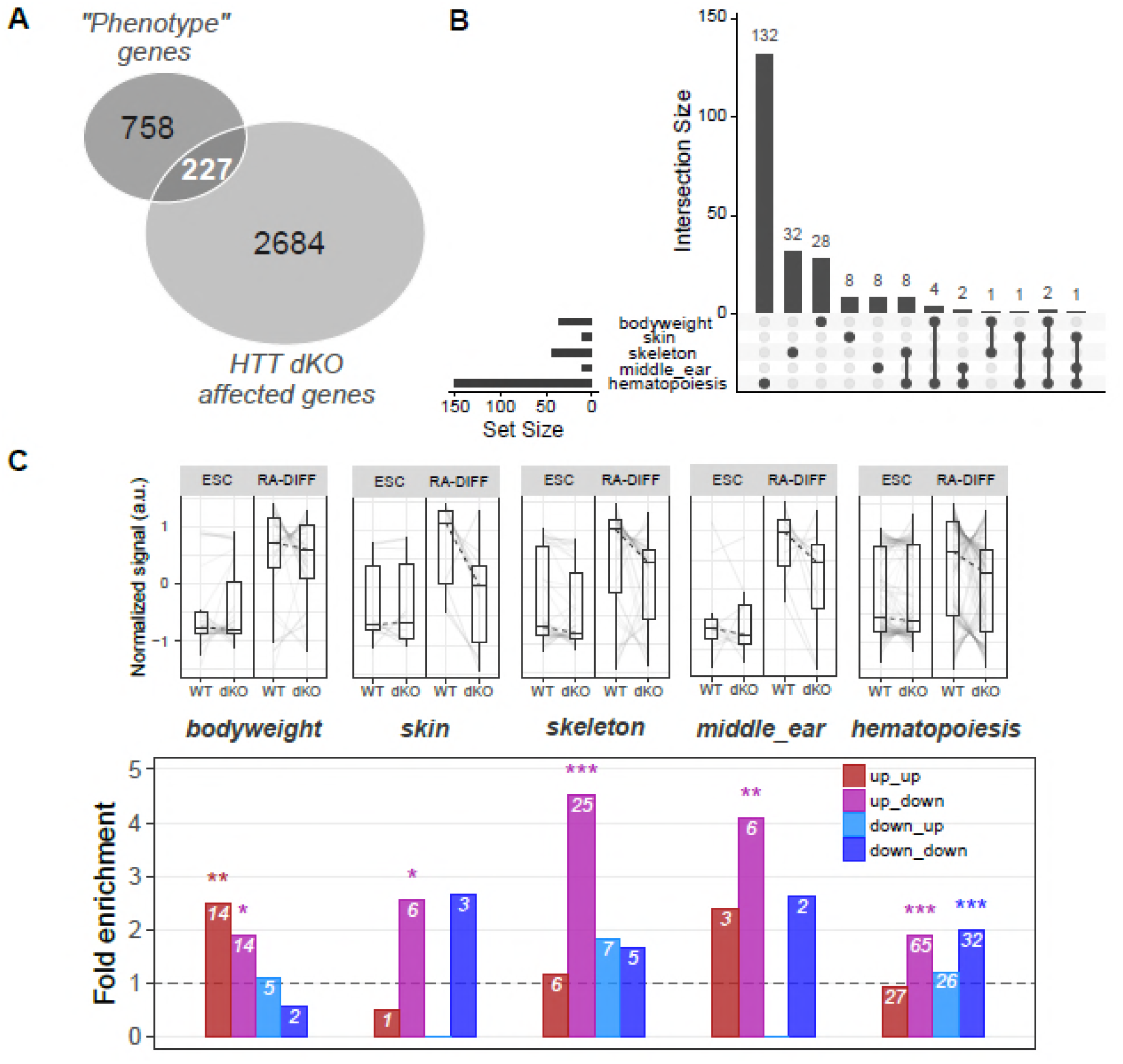
*Htt*^ex4/5^-null mutation, in the context of RA-differentiation, significantly perturbs genes implicated in ‘body weight’, ‘skin, skeleton, middle-ear development’ and ‘hematopoiesis’. A) Venn diagram representation of the intersection of genes relevant to *Htt* hypmorph related phenotypes (bodyweight, skin, skeleton, middle ear and hematopoiesis) and genes whose expression changes with *Htt*-genotype in stem cells (ESC) and during RA-EB differentiation (RA-Diff). The genes in this intersection were analyzed further in B) and C). B) The number of intersection genes associated with each of the *Htt-*hypomorph relevant phenotypes C) Expression trajectory plots (upper row) and enrichment analysis (lower row) of the four classes of genes (up-up; up-down; down-up; down-down; Fig. 6) in the intersection with developmental genes associated with *Htt* hypmorph related phenotypes.

### Integrative regulatory network analysis

Differentiation reflects the coordinated action of epigenetic regulators, transcription factors and post-transcriptional events among which those regulated by miRNAs on gene expression. Therefore, to determine whether *Htt*-inactivation may influence regulatory loops, thereby affecting the expression of downstream targets that may be involved in hypomophic mutation-associated mouse developmental phenotypes, we performed analysis with the above mentioned RA-differentiation-*Htt*-null gene sets to evaluate: 1) enrichment of targets of chromatin regulators in the ChEA experimental ChIP-seq database (https://www.ncbi.nlm.nih.gov/pubmed/20709693) and 2) potential miRNA regulators, by virtue of having a differentiation-*Htt* null expression pattern (down-down and down-up) that was opposite to that of their mRNA target genes (up-up and up-down) [25,26]. Consistent with previous reports, the results of the chromatin regulator analysis revealed a strong enrichment for targets of polycomb regulators (Fig. 8A) among the differentiation genes also regulated by *Htt* inactivating mutation, especially members of the PRC2 complex (Suppl. File S3). The second analysis revealed a set of 22 RA-differentiation-*Htt*-null responsive miRNAs whose known target genes’ expression pattern fulfilled our criteria (Suppl. File S1) (Fig. 8B-C, Suppl. File S4), with let-7b-5p and miR-329-3p the most significant for the down-down and down-up expression pattern classes, respectively. A functional enrichment analysis of the mRNAs regulated by our list of 22 miRNAs whose expression pattern fit our criteria (opposite to that of the binding miRNAs) confirmed enrichment of ‘up-up’ miRNA target genes in processes involved in ‘nervous system development, ‘axonogenesis’ and ‘synapse’, while the targets of ‘up-down’ miRNAs were enriched for terms related to ‘skeletal system development’, ‘organ morphogenesis’, ‘vasculature development’ ‘ear development’ and ‘skin development’ (Fig. 8B-C), implying a role for these regulators and target genes in the altered developmental phenotypes observed in mice with expression of the HD gene orthologue below a single functional allele.

**Figure 8.**
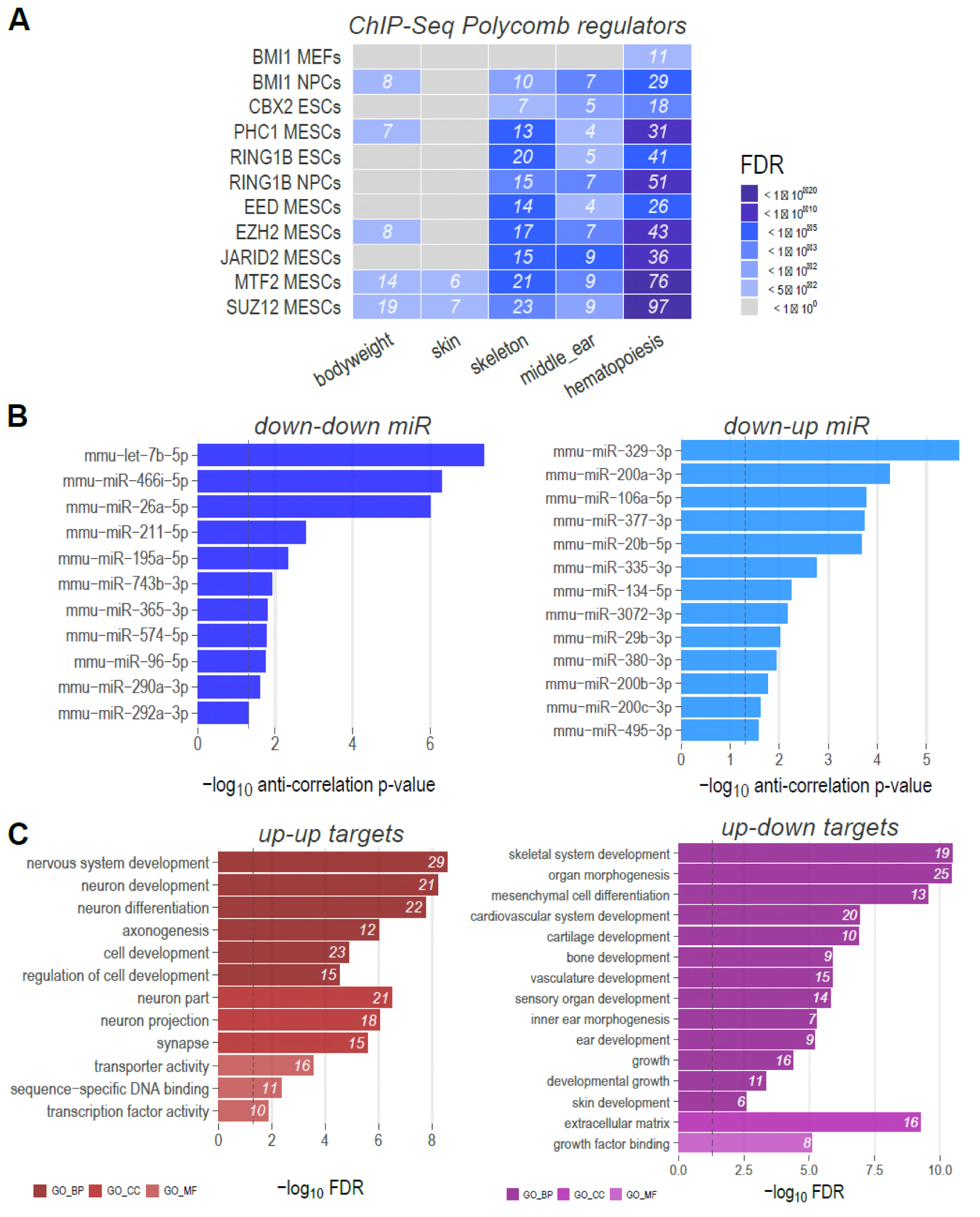
Regulatory network analysis highlights polycomb group protein and miRNAs as possible regulators. A) Enrichment analysis of genes whose expression is changed by *Hdh*^ex4/5/ex4/5^ null, that are involved in developmental phenotypes and regulated by Polycomb group proteins, based on ChEA ChIP-Seq annotation. Significance of enrichment is based on FDR values [color code as in the legend]. B) List of 22 miRNAs whose expression is altered by RA-differentiation and by *Hdh*^ex4/5^ null genotype, but whose change in expression is significantly anti-correlated relative to expression change observed in the miRNA target genes. miRNAs identified by these criteria are by definition either down-down or down-up. miRNAs are ranked according to the p-value associated with the anti-correlation of their targets. C) Functional enrichment analysis of mRNA regulated by the 22 miRNAs displayed in Panel B. Only mRNAs affected by RA-Diff and the *Htt* null mutation, as well as anti-correlated with respect to their miRNA regulators were selected for the analysis. Enrichment FDR values are displayed in the barplot.

## Discussion

HD is a neurodegenerative disorder for which there is as yet no disease-modifying intervention. It is hoped that approaches that decrease expression of the mutant gene, or both the mutant and normal gene, will provide benefit as suggested by observations from pre-clinical studies in model systems [27,28]. However, there is a need to better understand *HTT* function, in order to forecast potential effects of *HTT* silencing and lowering strategies and also to understand precisely how *HTT* CAG expansion mutation harms vulnerable cell populations that drive the features of the disease.

Our assessment of dosage effects, in which different hypomorphic mutations of the mouse HD gene orthologue attempt to rescue a null allele, has revealed critical involvement of *Htt* throughout development, from conception. First, the severe *Hdh*^neoQ111^ mutation, similarly to what previously shown for the homozygote null alleles, was characterize by a huntingtin level insufficient to properly initiate the organogenesis phase causing embryos to fail post-gastrulation, before head-folds become evident. Second, the intermediate *Hdh*^neoQ50^ mutation, did make sufficient huntingtin to overcome this first block, but this level was insufficient later, such that embryos failed after E10.5-12.5 [9], with survivors revealing several needful organ systems: brain, skeleton, middle and external ear, and skin barrier acquisition and fetal liver erythropoiesis. Third, the dose of huntingtin from the milder *Hdh*^neoQ20^ mutation was also wanting, but was sufficient for preventing most deficits observed with more severe mutations. Exceptions included the low penetrant external ear and sternum deficits and the enlarged ventricles noted previously [9,17] along with developmental delay after E14.5, although the small mice are born and can be viable (and are fertile) with assistance in the form of removal of normal littermates (with a wild-type allele) or hand-rearing [17]. These findings, taken together with previous observations that the cognate (neo-out) *Hdh*^Q20^, *Hdh*^50^, *Hdh*^Q111^ CAG repeat knock-in alleles (appropriately expressing endogenous mutant huntingtin with 20-, 50- and 111-glutamines) fully rescue the *Hdh*^ex4/5^ null [9,29] clearly demonstrate that the equivalent of a single functional HD orthologue allele (regardless of CAG repeat size) is needed for proper development. Our findings complement and enrich previous observations of aberrant external ear, vasculature, brain development and neurological deficits in mice that are compound heterozygote for neo-in hypomorphic alleles [17] and are also consistent with the report of a family segregating two *HTT* inactivating mutations, where compound heterozygotes exhibit decreased survival, global developmental delay and neurological deficits [15,16]. Indeed, our results strongly suggest that one or both of the *HTT* [4469+1G>A] and [8156T>A] mutations segregating in this family is functionally hypomorphic rather than completely inactivating.

Our molecular analysis of wild-type ESC and ESC with the *Hdh*^ex4/5^ null mutation, in a RA-morphogen-driven-paradigm [19-21], which we confirmed elicits broad effects on developmental genes, has nominated categories of regulators sensitive to huntingtin dosage that seem likely to play key roles in *Htt* function during organ system development. One important category is polycomb group proteins: PRC2 (Suz12, Eed, Jarid2 and Mtf2), implicated previously [12,13], as well as PRC1 (Bmi1, Ring1b), which are especially relevant for neurogenesis, hematopoiesis, skeleton and middle-ear formation [30-34]. The other major category is a group of miRNAs, among which Let-7b-5p and miR-129-3p, with known involvement in nervous system development, as well as skeletal, ear and skin development [35-37]. Precisely which processes these potential regulators orchestrate and how they may be disrupted by the lack of functional *Htt* in different organ systems remains to be investigated. However, several observations in severely hypomorphic embryos at different ages strongly suggest an impact at the level of progenitor stem cells implicating effectors of these regulators that are critical to cell adhesion or other aspects of cell-migration and can thereby influence lineage fate: 1) the pharyngeal arches were formed but the middle-ear fated Gsc-positive cells were located in the first arch, rather than in the second and third arches, implying delayed migration of these progenitor cells [38,39]; 2) ventral skin barrier formation was not permanently disrupted, only delayed at E18.5, suggesting aberrantly slow migration of epidermal precursors with delayed formation of a cornified layer and acquisition of barrier [40,41]; and 3) deficits in fetal liver hematopoiesis were transient, resolved by E18.5, consistent with delayed migration from the blood islands in the yolk sac, the site of embryonic hematopoiesis [42]. In addition, as reported previously [9,12,17,24,43], neurogenesis and brain development is abnormal in the absence of sufficient *Htt*. The results of our gene expression analysis support disruption of a process at the level of neural progenitor cells that influences the proportion of daughter neuronal and macroglial cell types, a process that normally involves the acquisition by radial glia of adherens junctions important for cell migration and proper specification, initially of neurons and later of astrocytes and oligodendrocytes [44].

It is evident that the inherited CAG repeat expansion mutation does not recapitulate the blatant effects produced by inheritance of two *HTT* inactivating mutations. HD mutation carriers, even homozygotes, are overtly indistinguishable from those who do not carry the mutation, until subtle changes presage the emergence of signs of the neurodegenerative disorder [45,46]. By contrast, though one active allele is sufficient [9], levels below a single allele’s worth of *HTT* function produce developmental consequences that our findings show can vary dramatically in scope and severity with dosage. Studies of decreased *Htt* dosage later in development in adults, rather than at conception, are more equivocal. Some reports show harmful consequences from neuron-specific knock-out, for example in [47], while a different strategy did not elicit harmful effects [48]. Our findings suggest that, if the harmful effects of the HD mutation are due to some opportunity for mischief that is provided by normal *HTT* function, then studies of the polycomb group protein genes and Let-7b-5p and miR-129-3p miRNAs as regulators of cellular adhesion may illuminate the biology that is disrupted by the HD mutation and, thereby, provide specific assays with which to evaluate therapeutics that aim to decrease or silence expression of the gene in individuals with the HD mutation.

## Methods

### Mice

All the mouse experiments were conducted in accordance with an IACUC approved protocol, through the MGH Subcommittee on Animal Research. The *Hdh*^ex4/5/ex4/5^ and *Hdh*^neoQ20^/*Hdh*^+^, *Hdh*^neoQ50^/*Hdh*^+^, *Hdh*^neoQ111^/*Hdh*^+^ alleles as well as the genotyping protocols have been described previously [6,7,9].

For skeletal preparations, embryos and newborns mice were eviscerated, fixed in ethanol, incubated in acetone and finally stained using alizarin red and alcian blue [49,50]. The *in vivo* epidermal barrier assay was performed on E18.5 embryos using a dye exclusion assay as reported[51]. Briefly, embryos were sacrificed, immersed in methanol solutions at different concentrations, then stained in 0.1% toluidine blue and washed several times with PBS again to remove the excess dye.

### Immunohistological analyses

Staining with hematoxylin and eosin (H&E) and immunostaining was performed by fixing frozen brain or skin sections [cryostat] (LEICA CM3050S), sectioned at 6 μm, in 4% Paraformaldehyde, incubating with primary antibodies at 4 °C overnight, while secondary antibody were used following vendor instructions. Anti-Proliferating Cell Nuclear Antigen (PCNA) was from Santa Cruz Biotechnology, Inc. Slides were rinsed and mounted using Vectashield with 4,6-diamidino-2-phenylindole (DAPI) (Vector Laboratories) for visualization of nuclei.

The DeadEnd™ Colorimetric TUNEL (TdT-mediated dUTP Nick-End Labeling) Assay (Promega) was used to detect apoptotic cells *in situ* accordingly to the vendor’s instructions.

### Protein extraction and immunoblot analysis

Total protein lysates were prepared by pulverizing skin tissue and extracted using RIPA (Boston Bio-Products) lysis buffer with protease inhibitor mixture (Roche). After Bradford protein assay (BIORAD), fifty micrograms subjected to 10% SDS–PAGE, transferred to nitrocellulose membranes (Schleicher and Schuell) and incubated with the following primary antibodies: anti-filaggrin (Covance), anti-loricrin (Covance) and anti-alpha tubulin antibody (Santa Cruz Biotechnology, Inc).

Mouse and rabbit secondary antibodies were used and the specific protein bands were detected using the ECL Plus kit (Pierce) and autoradiographic film (Hyperfilm ECL; Amersham Bioscience).

### Whole mount *in situ* hybridization

After dissection in PBS, embryos were fixed overnight in 4% paraformaldehyde at 4°C. RNA *in situ* hybridizations were performed as described previously (cold spring harbor lab method). Briefly, E11-11.5 mouse embryos were fixed overnight in 4% formaldehyde in PBS, then dehydrated in methanol and stored at −20°C until use. The digoxigenin-labelled RNA probes were used at 0.5 ug/ml. Alkaline phosphatase-conjugated anti-digoxigenin Fab fragments were used at 1:5000. Color reactions were carried out over time periods ranging from 2 hours to overnight. Embryos were mounted in 80% glycerol before being photographed. Three to four embryos were evaluated for each marker.

### Flow cytometry

Fetal liver and fetal thymus were harvested from E14.5 and E18.5 *Hdh*^+^/*Hdh*^+^ and *Hdh*^neo-^ inQ50/*Hdh*^ex4/5^ embryos and single cell suspensions were obtained by mechanical disruption. Fetal liver cells were stained with Ter119, CD71, Mac-1, Gr-1 B220 and CD19, and thymocytes were stained with CD4 and CD8a cell surface markers, respectively. Flow cytometry was performed on a two-laser FACSCanto (BD Biosciences). FCS files were analyzed by FlowJo software (Tree Star). Antibodies used were CD4 (L3T4), CD8a (53-6.7), CD19 (1D3), Mac-1 (M1/70), B220 (RA3-6B2), Gr-1 (RB6-8C5), Ter119, CD71 (C2) (BD Pharmingen, eBioscience).

### Colony forming cell (CFC) assays

Methylcellulose colony forming cell (CFC) assays was performed on fetal liver cells of E14.5 and E18.5 *Hdh*^+^/*Hdh*^+^ and *Hdh*^neo-inQ50^/*Hdh*^ex4/5^ mice as previously described (Yoshida et al G&D 2008). Briefly, 5 × 10^4^ cells were cultured with Methocult M3434 (Stem Cell Technologies) supplemented with hTPO (50 ng/ml) in 35-mm culture dishes (NUNC 174926) in duplicates. Colonies were scored from day 2 to day 17 according to the technical manual and previously described criteria [52], and were confirmed by May-Giemsa staining (Harleco) of cytospins (400 rpm, 5 min) of individual colonies. Cytokines were purchased from R&D Systems.

### Cell culture

Wild-type and huntingtin null *Hdh*^ex4/5/ex4/5^ mouse embryonic stem cell lines were described previously [9,12,53]. Differentiation was performed essentially as described in Bibel et al., 2007 [54]. In brief, ESCs were deprived of feeder cells for 4 passages, then 3 x (10^6^) cells were used for formation of embryoid bodies (EBs). EBs were grown in non-adherent bacterial dishes (Greiner, Germany) for 8 days. Retinoic acid (5 uM, SIGMA) was added from day 4 to day 8 and medium was changed every other day. Subsequently, EBs were dissociated by trypsin digestion and plated on Poly-Ornithine (SIGMA) and laminin (SIGMA) coated plates to obtain RA-differentiated cells (RA-DIFF). Two hours after plating, RA-differentiated cells were collected for different analyses.

### RNA Isolation and RNAseq, miRNAseq library preparation and sequencing

RNA was extracted from cell lines by using TRIzol reagent (Life technologies), following manufacturer’s instructions. All RNA samples were subjected to DNAse I treatment (Ambion). RNA sequencing was performed following the protocol described by the Broad Institute (Cambridge, MA)[55,56]. Briefly, poly-A-plus mRNA, was retro-transcribe to cDNA using a strand specific dUTP method, random hexamers and amplified by PCR using bar-coded DNA adaptors from Illumina. HiSeq2000 platform and 50bp pair-end (PE) was used for sequencing obtaining 50-75M reads/library.

Small RNA library preparation (mainly miRNAs) was obtained using the Illumina TruSeq Small RNA protocol (Illumina) according to manufacturer instructions and libraries were sequenced using single-end, 50bp reads on HiSeq2000 platform, obtaining 45-90M reads/library.

### RNA-Sequencing and miRNA-Sequencing data analysis

For RNA-Seq, 50 bp paired-end reads were were aligned to the mouse genome (GRCm38.p4) with Tophat (version 2.0.14, default settings), using the Gencode M6 transcript annotation as transcriptome guide. All programs were used with default settings unless otherwise specified. Mapped reads (Suppl. Table 1) were assembled into transcripts guided by reference annotation (Gencode M6) with Cufflinks (version 2.2.1). Expression levels were quantified by HTSeq (Version 0.6.1, --mode intersection-nonempty) using Gencode M6 annotation. Normalization was performed using the TMM method implemented in edgeR. Differential expression was calculated using edgeR (dispersion 0.1, pval < 0.05, log2 fc >0.75, log2CPM > 0).

For miRNA-Seq, 50 bp single-end reads were trimmed against Illumina’s TruSeq Small RNA 3’ (TGGAATTCTCGGGTGCCAAGG) and 5’ (GUUCAGAGUUCUACAGUCCGACGAUC) adapters using cutadapt(v. 1.3) with parameters –e 0.1 –O 5 –m 15[57]. Next, trimmed sequence reads with length of 16-25bp were mapped to 1915 known mature miRNA sequences from miRBase (release 21) database [58], using BWA aln (v. 0.7.5a-r418) with parameter –n 1 [59]. Uniquely mapped reads (Suppl. Table 1) were identified by selecting alignments with non-zero mapping quality and “XT:A:U” tag using samtools (v 0.1.18)[60]. Normalization was performed using the TMM method (BWA and TMM were suggested previously[61]). Differential expression was calculated using edgeR (dispersion 0.1, pval < 0.05, log2 fc >0.75, log2CPM > 0)[62].

### Computational analyses

Functional annotation of gene lists and enrichment analysis with Gene Ontology terms and KEGG or REACTOME pathways were performed with the clusterProfiler Bioconductor package. Neuron, astrocyte and oligodendrocyte markers were downloaded from Cahoy et al., 2008 [23], while lists of genes implicated in the different phenotypes were manually created based on multiple papers/web-resources [39, 63-68] (Suppl. Table 2). The significance of custom enrichments was measured with one-sided Fisher exact test.

Collections of murine miRNA targets were downloaded from miRTarBase (Release 6.1) [69]. For each miRNA – target mRNA couple, Pearson’s correlation values were calculated from standard normalized expression values. For each miRNA, the negative shift in the distribution of target correlation values was measured with one-sample one-sided Wilcoxon test (significance threshold: P < 0.05).

## Acknowledgments

We thank Jolene Guide for technical support with mouse breeding and Ilaria Sanvido for assisting in the ES cell culture experiments. This work was supported by the National Institutes of Health (USA) (grant NS32765), the CHDI Foundation Inc. (MEM, JFG), an Anonymous Donor (MEM) and the University of Trento (MB). MB was a recipient of a Marie Skłodowska-Curie reintegration fellowship (the European Union’s Horizon 2020 research and innovation programme) under the grant agreement No. 706567.

---

**Suppl. Figure 1.**
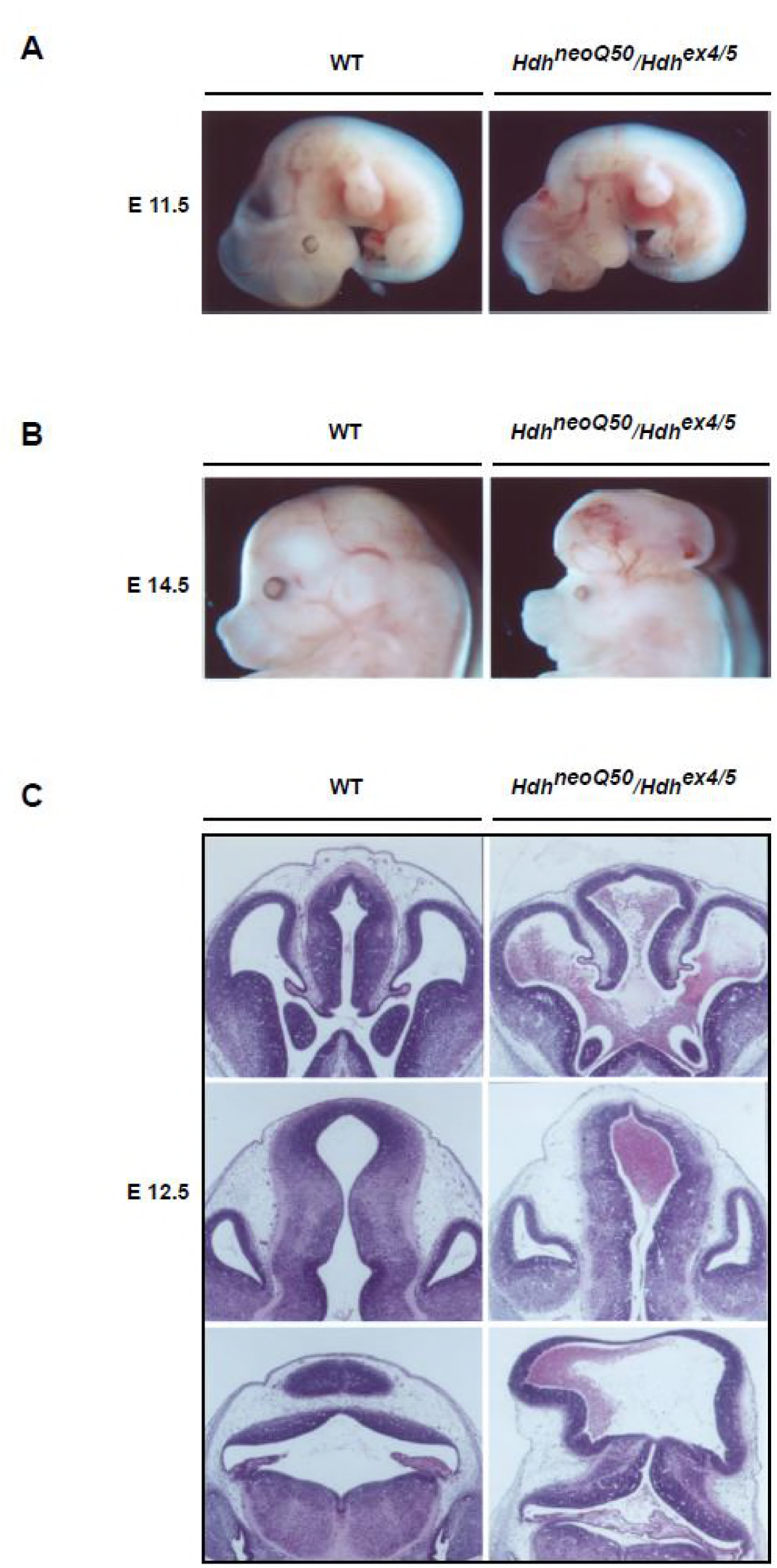
Morphological characterization of brain vasculature defects in *Hdh* hypomorhic mice. A-B) Representative pictures of WT and *Hdh*^neoQ50^/*Hdh*^ex4/5^ hypomorphic embryos at E 11.5 and E 14.5 developmental stages. Visible vasculature defects are present in hypomorphic embryos, especially at the level of the brain. C) Representative pictures of histological sections of *Hdh*^neoQ50^/*Hdh*^ex4/5^ hypomorphic embryos brains confirm enlarged ventricles with extensive blood accumulation.

**Suppl. Figure 2.**
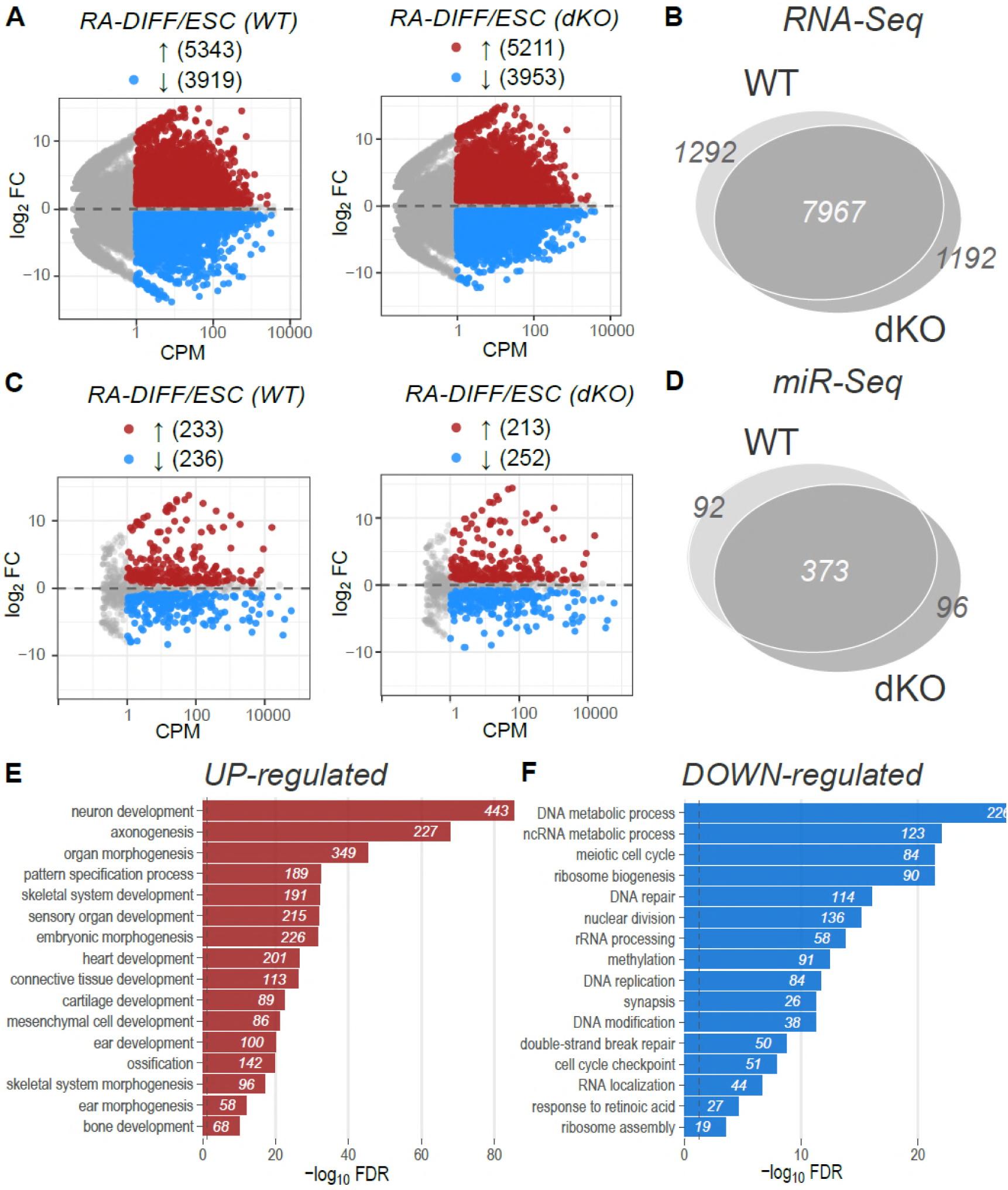
The transcriptional changes induced by RA differentiation are quantitatively and qualitatively similar in *Htt* wild-type and *Hdh*^ex4/5/ex4/5^ cells. A) M (log ratio) and A (mean average) (MA) plot representations of mRNA-seq pairwise comparisons of RA-Diff cells by wild-type (WT) or *Hdh*^ex4/5/ex4/5^ *Htt* null (dKO) genotypes showing the average log10 signal (Counts Per Million - CPM) against the log2 Fold Change (FC) for each gene. Genes significantly up-regulated or down-regulated in the comparison are highlighted in red and blue, respectively. Numbers of differentially expressed genes are displayed (parenthesis). B) The Venn diagram reports the total number of genes that are commonly or specifically dysregulated during differentiation (transition from ESC to RA-DIFF), comparing cells with *Htt* wild-type (WT) or *Htt-null (*dKO) genotypes. C) MA plot representations of miRNA-seq pairwise comparisons. Legends, abbreviation and colors as in A). D) The Venn diagram reports the total number of miRNAs that are commonly or specifically changed during RA differentiation by wild-type (WT) or *Hdh*^ex4/5/ex4/5^ *Htt* null (dKO) genotypes. Panel E) Cluster 1 contains genes that are highly expressed in ESC and whose expression decreases during RA differentiation. Bar plots for this cluster report the most enriched GO-terms describing biological processes. The number of genes within each GO-term is also indicated (number within bars). F) Cluster 2 groups genes that are poorly expressed in ESC, but strongly upregulated during RA differentiation. Bar plots report the 5 most enriched GO terms associated with genes in Cluster 2. The number of genes within each GO-term is also indicated (number within bars). Genes belonging to Cluster 1 and 2 show similar behavior in cells with *Htt* wild-type (WT) or *Htt*-null (dKO) genotypes.

**Suppl. Figure 3.**
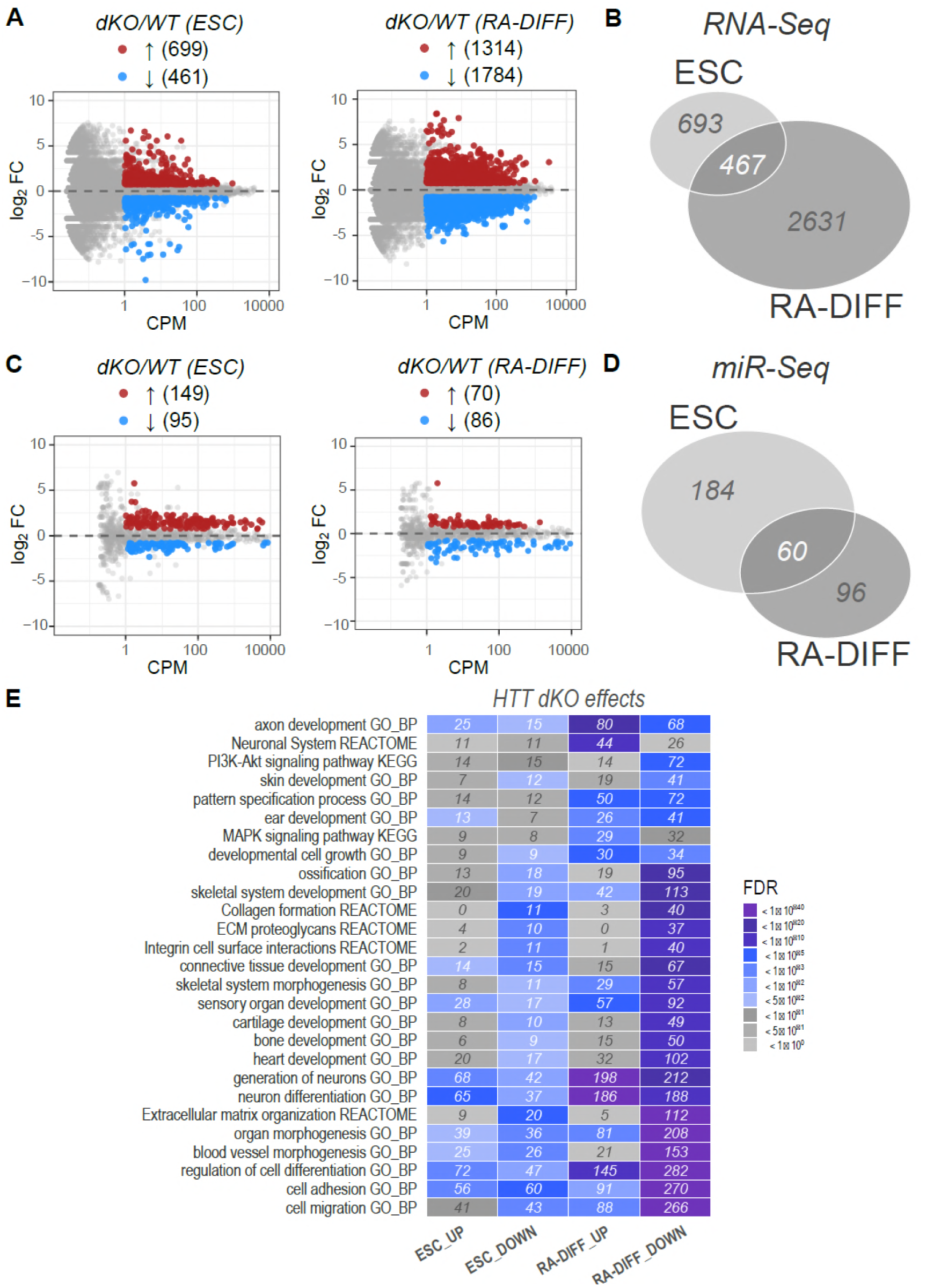
The transcriptional changes due to *Hdh*^ex4/5/ex4/5^ null genotype are quantitatively and qualitatively different in ESC and RA-DIFF cells and are milder compared to changes by RA differentiation. A) M (log ratio) and A (mean average) (MA) plot representations of mRNA-seq pairwise comparisons of *Hdh*^ex4/5/ex4/5^ *Htt* null (dKO) versus wild-type (WT) genotypes in ESC and RA-diff cells showing the average log10 signal (Counts Per Million - CPM) against the log2 Fold Change (FC) for each gene. Genes significantly up-regulated or down-regulated in the comparison are highlighted in red and blue, respectively. Numbers of differentially expressed genes are displayed (parenthesis). B) The Venn diagram reports the total number of genes that are commonly or specifically dysregulated comparing cells with *Htt* wild-type (WT) or *Htt*-null (dKO) genotypes in ESC and RA-diff cells. C) MA plot representations of miRNA-seq pairwise comparisons. Legends, abbreviation and colors as in A). D) The Venn diagram reports the total number of miRNAs that are commonly or specifically dysregulated comparing cells with *Htt* wild-type (WT) or *Htt*-null (dKO) genotypes in ESC and RA-diff cells. E) Heatmap reports the top 25 most enriched Reactome and KEGG pathways associated with genes up and down-regulated in ESC and RA-DIFF cells in absence of huntingtin. The number of affected genes within each pathway is indicated (numbers in the cells).

**Suppl. Figure 4.**
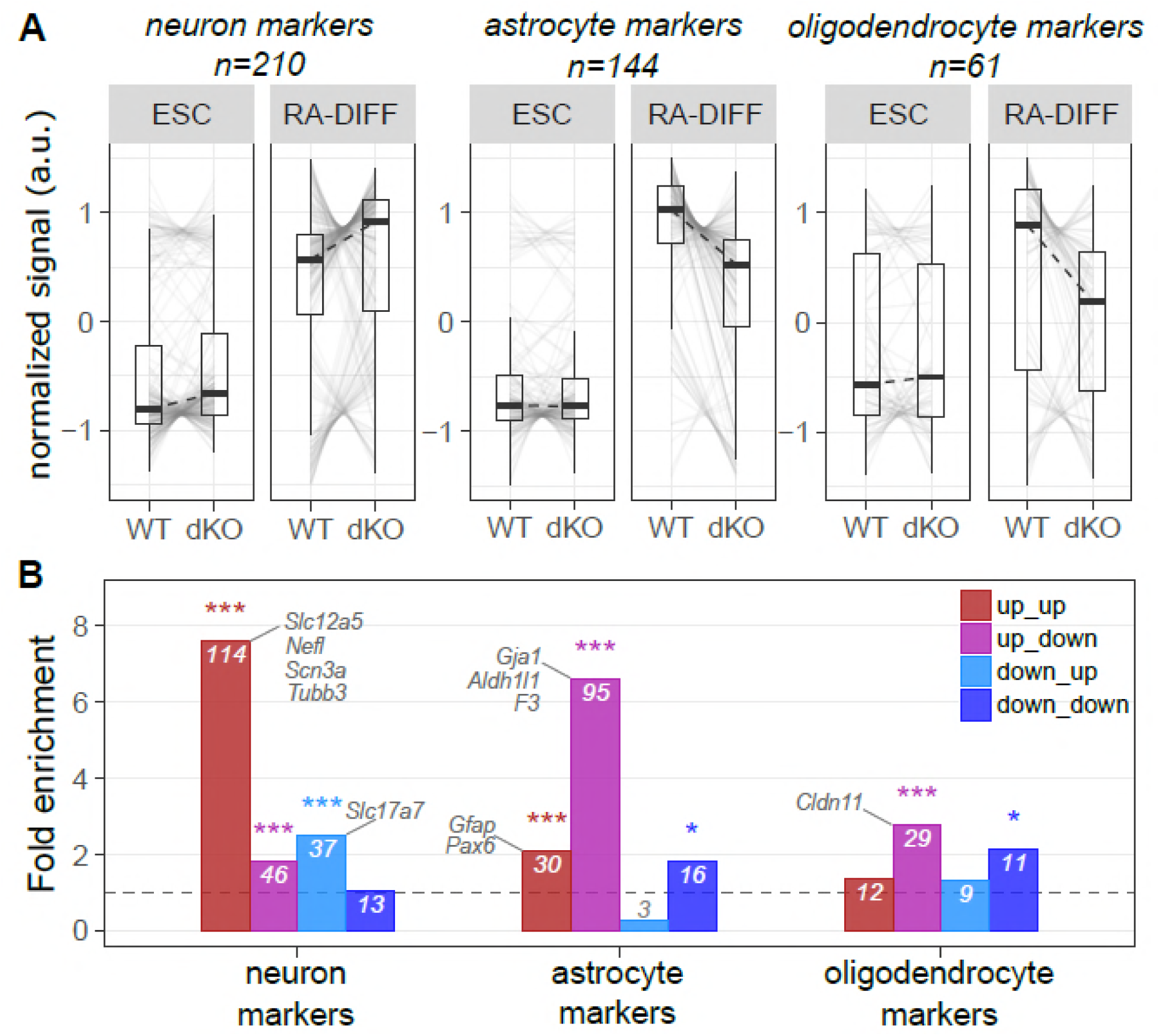
*Hdh*^ex4/5/ex4/5^ null mutation alters expression of genes involved in neuron-glial specification. A) Expression trajectory plots describing variations in transcriptional levels of genes involved in neuron-glial specification during RA-DIFF: transcriptional changes for neuron markers, astrocyte markers, oligodendrocyte markers[23] between the two *Htt* genotypes (wild-type; WT and *Htt*-null; dKO) and two developmental stages (ESC and RA-DIFF) are shown. The number of genes defining each class of markers is indicated at the top of each plot. B) Enrichment bar plots depicting the enrichment values of neuron, astrocyte and oligodendrocyte markers among the 4 gene classes (up_up; up_down; down_down; down_up, as described in Fig. 3A) of genes affected by RNA-differentiation and by *Hdh*^ex4/5/ex4/5^ null mutation. The number of genes enriched in each of the 4 classes is indicated within each column. Statistical significance, measured by Fisher exact test is indicated by asterisks: (*) P<0.05, (***) P<0.01.

**Suppl. Table 1**

Table depicted total number of raw and aligned reads for RNA-seq (A) and miRNA-seq (B) experiments. Data are related to Figure 6 and Suppl. Fig 2-3.

**Suppl. Table 2**

Table summarized the source of the papers/webtools used to create the manually annotated genes’ lists for genes-associated-phenotypes. Data are related to Figure 7 of the main text.

**Suppl. File 1**

The excel file presented two spreadsheet describing RNA-seq and miRNA-seq full data. For each comparison a color code was used to highlight genes/miRNAs that were significantly downregulated (red) or upregulated (green). Log_2_ CPM or Log_2_Fold Change (FC) values were obtained for each gene or miRNA. RNA-seq: Gene symbol and Ensembl ID, gene type (protein coding or non-coding), full description of the gene name, chromosomal and strand location as well as genomic coordinates were described. For each gene its fitting to a functional enrichment class [up-up, up-down, down-down, down-up] (see also Fig. 6) or its role as neuron, astrocyte, oligodendrocyte_marker (see also Suppl. Fig. 4) was reported, while its contribution to developmental phenotypes (middle_ear_development, hematopoiesis, skeleton, skin, bodyweight) was presented in columns I-Q.

miRNA-seq: Similarly to what presented for genes, for each miRNA its belonging to the functional enrichment class [up-up, up-down, down-down, down-up] (see also Fig. 8) was reported, while its previous correlation with HD mutation [Hoss_2015, Hoss_2014, Lee_2011_1, Lee_2011_2, Marti_2010_1, Marti_2010_2] was assessed as described in columns C-J.

**Suppl. File 2**

The excel file presented spreadsheets describing significant GO terms and pathways (Reactome-Keggs) for the 10 different comparisons presented in Fig. 6 – 8 [differentiation_up, differentiation_down, *Htt*dKO_ESC_up, *Htt*dKO_ESC_down, *Htt*dKO_RA-DIFF_up, *Htt*dKO_RA-DIFF_down, up-up, up-down, down-up, down-down). For each comparison/spreadsheet the ontology ID and description, fold enrichment, P-value, False Discovery rate (FDR) significance and annotated genes were reported.

**Suppl. File 3**

The excel file reported the ChEA analysis related to Fig. 8. For each ChEA datasets, the phenotype enrichment (middle_ear_development, hematopoiesis, skeleton, skin, bodyweight), combined score, fold enrichment, p_value, False Discovery Rate (FDR) significance and the annotated genes were described.

**Suppl. File 4**

The excel file reported 4 different spreadsheets describing i. miRNAs with anticorrelated target, ii. miRNAs target interactions, iii. enrichment analysis for UP_UP miRNAs targets and iv. enrichment analysis for UP_DOWN miRNAs targets to support results presented in Fig. 8. For table describing miRNAs with anti-correlated targets, the miRNA ID, miR_class (down_enhanced, down_counteracted), number of miRNAs annotated and anti-correlated target genes, average Spearman and Pearson’s correlation and pvalue were reported. Similarly, for miRNAs target interactions, the mRNA target ID and description, technical evidence of miRNA-target interaction as well as its belonging to the developmental phenotypes (middle_ear_development, hematopoiesis, skeleton, skin, bodyweight) were described. Finally enrichment analysis for miRNAs targets, presenting ontology ID and description, fold enrichment, P-value, False Discovery rate (FDR) significance and annotated genes were reported for UP_UP and UP_DOWN classes (see Fig. 8 and results).

## References

1. Harper PS (1999) Huntington’s disease: a clinical, genetic and molecular model for polyglutamine repeat disorders. Philos Trans R Soc Lond B Biol Sci 354: 957–961.

2. (1993) A novel gene containing a trinucleotide repeat that is expanded and unstable on Huntington’s disease chromosomes. The Huntington’s Disease Collaborative Research Group. Cell 72: 971–983.

3. Genetic Modifiers of Huntington’s Disease C (2015) Identification of Genetic Factors that Modify Clinical Onset of Huntington’s Disease. Cell 162: 516–526.

4. Gusella JF, MacDonald ME (2006) Huntington’s disease: seeing the pathogenic process through a genetic lens. Trends Biochem Sci 31: 533–540.

5. Lee JM, Ramos EM, Lee JH, Gillis T, Mysore JS, et al. (2012) CAG repeat expansion in Huntington disease determines age at onset in a fully dominant fashion. Neurology 78: 690–695.

6. Ambrose CM, Duyao MP, Barnes G, Bates GP, Lin CS, et al. (1994) Structure and expression of the Huntington’s disease gene: evidence against simple inactivation due to an expanded CAG repeat. Somat Cell Mol Genet 20: 27–38.

7. Duyao MP, Auerbach AB, Ryan A, Persichetti F, Barnes GT, et al. (1995) Inactivation of the mouse Huntington’s disease gene homolog Hdh. Science 269: 407–410.

8. Wheeler VC, Auerbach W, White JK, Srinidhi J, Auerbach A, et al. (1999) Length-dependent gametic CAG repeat instability in the Huntington’s disease knock-in mouse. Hum Mol Genet 8: 115–122.

9. White JK, Auerbach W, Duyao MP, Vonsattel JP, Gusella JF, et al. (1997) Huntingtin is required for neurogenesis and is not impaired by the Huntington’s disease CAG expansion. Nat Genet 17: 404–410.

10. Zeitlin S, Liu JP, Chapman DL, Papaioannou VE, Efstratiadis A (1995) Increased apoptosis and early embryonic lethality in mice nullizygous for the Huntington’s disease gene homologue. Nat Genet 11: 155–163.

11. Nasir J, Floresco SB, O’Kusky JR, Diewert VM, Richman JM, et al. (1995) Targeted disruption of the Huntington’s disease gene results in embryonic lethality and behavioral and morphological changes in heterozygotes. Cell 81: 811–823.

12. Biagioli M, Ferrari F, Mendenhall EM, Zhang Y, Erdin S, et al. (2015) Htt CAG repeat expansion confers pleiotropic gains of mutant huntingtin function in chromatin regulation. Hum Mol Genet 24: 2442–2457.

13. Seong IS, Woda JM, Song JJ, Lloret A, Abeyrathne PD, et al. (2010) Huntingtin facilitates polycomb repressive complex 2. Hum Mol Genet 19: 573–583.

14. Woda JM, Calzonetti T, Hilditch-Maguire P, Duyao MP, Conlon RA, et al. (2005) Inactivation of the Huntington’s disease gene (Hdh) impairs anterior streak formation and early patterning of the mouse embryo. BMC Dev Biol 5: 17.

15. Lopes F, Barbosa M, Ameur A, Soares G, de Sa J, et al. (2016) Identification of novel genetic causes of Rett syndrome-like phenotypes. J Med Genet 53: 190–199.

16. Rodan LH, Cohen J, Fatemi A, Gillis T, Lucente D, et al. (2016) A novel neurodevelopmental disorder associated with compound heterozygous variants in the huntingtin gene. Eur J Hum Genet 24: 1826–1827.

17. Auerbach W, Hurlbert MS, Hilditch-Maguire P, Wadghiri YZ, Wheeler VC, et al. (2001) The HD mutation causes progressive lethal neurological disease in mice expressing reduced levels of huntingtin. Hum Mol Genet 10: 2515–2523.

18. Presland RB, Dale BA (2000) Epithelial structural proteins of the skin and oral cavity: function in health and disease. Crit Rev Oral Biol Med 11: 383–408.

19. Duester G (2008) Retinoic acid synthesis and signaling during early organogenesis. Cell 134: 921–931.

20. Niederreither K, Dolle P (2008) Retinoic acid in development: towards an integrated view. Nat Rev Genet 9: 541–553.

21. Rhinn M, Dolle P (2012) Retinoic acid signalling during development. Development 139: 843–858.

22. Yu G, Wang LG, Han Y, He QY (2012) clusterProfiler: an R package for comparing biological themes among gene clusters. OMICS 16: 284–287.

23. Cahoy JD, Emery B, Kaushal A, Foo LC, Zamanian JL, et al. (2008) A transcriptome database for astrocytes, neurons, and oligodendrocytes: a new resource for understanding brain development and function. J Neurosci 28: 264–278.

24. Conforti P, Camnasio S, Mutti C, Valenza M, Thompson M, et al. (2013) Lack of huntingtin promotes neural stem cells differentiation into glial cells while neurons expressing huntingtin with expanded polyglutamine tracts undergo cell death. Neurobiol Dis 50: 160–170.

25. Huang JC, Babak T, Corson TW, Chua G, Khan S, et al. (2007) Using expression profiling data to identify human microRNA targets. Nat Methods 4: 1045–1049.

26. Lim LP, Lau NC, Garrett-Engele P, Grimson A, Schelter JM, et al. (2005) Microarray analysis shows that some microRNAs downregulate large numbers of target mRNAs. Nature 433: 769–773.

27. Tabrizi S, Leavitt B, Kordasiewicz H, Czech C, Swayze E, et al. (2018) Effects of IONIS-HTTRx in Patients with Early Huntington’s Disease, Results of the First HTT-Lowering Drug Trial (CT.002). Neurology 90: CT.002.

28. Rodrigues FB, Wild EJ (2018) Huntington’s Disease Clinical Trials Corner: February 2018. J Huntingtons Dis 7: 89–98.

29. Wheeler VC, Gutekunst CA, Vrbanac V, Lebel LA, Schilling G, et al. (2002) Early phenotypes that presage late-onset neurodegenerative disease allow testing of modifiers in Hdh CAG knock-in mice. Hum Mol Genet 11: 633–640.

30. O’Carroll D, Erhardt S, Pagani M, Barton SC, Surani MA, et al. (2001) The polycomb-group gene Ezh2 is required for early mouse development. Mol Cell Biol 21: 4330–4336.

31. Schumacher A, Faust C, Magnuson T (1996) Positional cloning of a global regulator of anterior-posterior patterning in mice. Nature 384: 648.

32. van der Lugt NM, Domen J, Linders K, van Roon M, Robanus-Maandag E, et al. (1994) Posterior transformation, neurological abnormalities, and severe hematopoietic defects in mice with a targeted deletion of the bmi-1 proto-oncogene. Genes Dev 8: 757–769.

33. Ding X, Lin Q, Ensenat-Waser R, Rose-John S, Zenke M (2012) Polycomb group protein Bmi1 promotes hematopoietic cell development from embryonic stem cells. Stem Cells Dev 21: 121–132.

34. Molofsky AV, Pardal R, Iwashita T, Park IK, Clarke MF, et al. (2003) Bmi-1 dependence distinguishes neural stem cell self-renewal from progenitor proliferation. Nature 425: 962–967.

35. Wu J, Qian J, Li C, Kwok L, Cheng F, et al. (2010) miR-129 regulates cell proliferation by downregulating Cdk6 expression. Cell Cycle 9: 1809–1818.

36. Gokbuget D, Pereira JA, Bachofner S, Marchais A, Ciaudo C, et al. (2015) The Lin28/let-7 axis is critical for myelination in the peripheral nervous system. Nat Commun 6: 8584.

37. Zhang Y, Yin B, Shu B, Liu Z, Ding H, et al. (2017) Differential expression of microRNA let-7b-5p regulates burn-induced hyperglycemia. Oncotarget 8: 72886–72892.

38. Rivera-Perez JA, Wakamiya M, Behringer RR (1999) Goosecoid acts cell autonomously in mesenchyme-derived tissues during craniofacial development. Development 126: 3811–3821.

39. Mallo M (2001) Formation of the middle ear: recent progress on the developmental and molecular mechanisms. Dev Biol 231: 410–419.

40. Li Q, Lu Q, Hwang JY, Buscher D, Lee KF, et al. (1999) IKK1-deficient mice exhibit abnormal development of skin and skeleton. Genes Dev 13: 1322–1328.

41. Kashiwagi M, Morgan BA, Georgopoulos K (2007) The chromatin remodeler Mi-2beta is required for establishment of the basal epidermis and normal differentiation of its progeny. Development 134: 1571–1582.

42. Keller G, Lacaud G, Robertson S (1999) Development of the hematopoietic system in the mouse. Exp Hematol 27: 777–787.

43. Yu MS, Tanese N (2017) Huntingtin Is Required for Neural But Not Cardiac/Pancreatic Progenitor Differentiation of Mouse Embryonic Stem Cells In vitro. Front Cell Neurosci 11: 33.

44. Martynoga B, Drechsel D, Guillemot F (2012) Molecular control of neurogenesis: a view from the mammalian cerebral cortex. Cold Spring Harb Perspect Biol 4.

45. Lee JK, Mathews K, Schlaggar B, Perlmutter J, Paulsen JS, et al. (2012) Measures of growth in children at risk for Huntington disease. Neurology 79: 668–674.

46. Paulsen JS, Nopoulos PC, Aylward E, Ross CA, Johnson H, et al. (2010) Striatal and white matter predictors of estimated diagnosis for Huntington disease. Brain Res Bull 82: 201–207.

47. Dietrich P, Johnson IM, Alli S, Dragatsis I (2017) Elimination of huntingtin in the adult mouse leads to progressive behavioral deficits, bilateral thalamic calcification, and altered brain iron homeostasis. PLoS Genet 13: e1006846.

48. Wang G, Liu X, Gaertig MA, Li S, Li XJ (2016) Ablation of huntingtin in adult neurons is nondeleterious but its depletion in young mice causes acute pancreatitis. Proc Natl Acad Sci U S A 113: 3359–3364.

49. Gruss P, Kessel M (1991) Axial specification in higher vertebrates. Curr Opin Genet Dev 1: 204–210.

50. Kessel M (1992) Respecification of vertebral identities by retinoic acid. Development 115: 487–501.

51. Byrne C, Avilion AA, O’Shaughnessy RF, Welti JC, Hardman MJ, editors (2010) Whole-Mount Assays for Gene Induction and Barrier Formation in the Developing Epidermis.: Humana Press, Totowa, NJ.

52. Nakahata T, Ogawa M (1982) Hemopoietic colony-forming cells in umbilical cord blood with extensive capability to generate mono- and multipotential hemopoietic progenitors. J Clin Invest 70: 1324–1328.

53. Jacobsen JC, Gregory GC, Woda JM, Thompson MN, Coser KR, et al. (2011) HD CAG-correlated gene expression changes support a simple dominant gain of function. Hum Mol Genet 20: 2846–2860.

54. Bibel M, Richter J, Lacroix E, Barde YA (2007) Generation of a defined and uniform population of CNS progenitors and neurons from mouse embryonic stem cells. Nat Protoc 2: 1034–1043.

55. Sugathan A, Biagioli M, Golzio C, Erdin S, Blumenthal I, et al. (2014) CHD8 regulates neurodevelopmental pathways associated with autism spectrum disorder in neural progenitors. Proc Natl Acad Sci U S A 111: E4468–4477.

56. Levin JZ, Yassour M, Adiconis X, Nusbaum C, Thompson DA, et al. (2010) Comprehensive comparative analysis of strand-specific RNA sequencing methods. Nat Methods 7: 709–715.

57. Martin M (2011) Cutadapt removes adapter sequences from high-throughput sequencing reads. EMBnetjournal 17: 10–12.

58. Kozomara A, Griffiths-Jones S (2014) miRBase: annotating high confidence microRNAs using deep sequencing data. Nucleic Acids Res 42: D68–73.

59. Li H, Durbin R (2010) Fast and accurate long-read alignment with Burrows-Wheeler transform. Bioinformatics 26: 589–595.

60. Li H, Handsaker B, Wysoker A, Fennell T, Ruan J, et al. (2009) The Sequence Alignment/Map format and SAMtools. Bioinformatics 25: 2078–2079.

61. Tam S, Tsao MS, McPherson JD (2015) Optimization of miRNA-seq data preprocessing. Brief Bioinform 16: 950–963.

62. Robinson MD, McCarthy DJ, Smyth GK (2010) edgeR: a Bioconductor package for differential expression analysis of digital gene expression data. Bioinformatics 26: 139–140.

63. Kemper KE, Visscher PM, Goddard ME (2012) Genetic architecture of body size in mammals. Genome Biol 13: 244.

64. Jung HJ, Tatar A, Tu Y, Nobumori C, Yang SH, et al. (2014) An absence of nuclear lamins in keratinocytes leads to ichthyosis, defective epidermal barrier function, and intrusion of nuclear membranes and endoplasmic reticulum into the nuclear chromatin. Mol Cell Biol 34: 4534–4544.

65. Richard G (2004) Molecular genetics of the ichthyoses. Am J Med Genet C Semin Med Genet 131C: 32–44.

66. Li L, Liu B, Wapinski OL, Tsai MC, Qu K, et al. (2013) Targeted disruption of Hotair leads to homeotic transformation and gene derepression. Cell Rep 5: 3–12.

67. Chen C, Jiang Y, Xu C, Liu X, Hu L, et al. (2016) Skeleton Genetics: a comprehensive database for genes and mutations related to genetic skeletal disorders. Database (Oxford) 2016.

68. Mallo M (1998) Embryological and genetic aspects of middle ear development. Int J Dev Biol 42: 11–22.

69. Chou CH, Chang NW, Shrestha S, Hsu SD, Lin YL, et al. (2016) miRTarBase 2016: updates to the experimentally validated miRNA-target interactions database. Nucleic Acids Res 44: D239–247.

